# Modularity-dependent storage of dynamic spiking patterns: bridging micro- and mesoscopic representations

**DOI:** 10.64898/2026.02.02.703247

**Authors:** Marianna Angiolelli, Antonio De Candia, Pierpaolo Sorrentino, Simonetta Filippi, Letizia Chiodo, Christian Cherubini, Silvia Scarpetta

## Abstract

Biological systems rely on asynchronous and temporally overlapping dynamics, allowing for the concurrent activation of multiple processes. This principle is particularly evident in brain function, where cognitive tasks engage distributed, interacting regions rather than sequentially isolated ones. To investigate the mechanisms enabling such coordination, we study a modular spiking neural network composed of leaky integrate-and-fire neurons and governed by spike-timing-dependent plasticity (STDP). Our model stores modular spatiotemporal patterns both at the mesoscopic level (sequences of modules) and at the microscopic level (precise spike timings) and includes a parameter, *η*, which regulates the degree of temporal overlap between modules’ activations. By tuning *η*, the network transitions from sequential to overlapping regimes, ranging from synfire chain-like dynamics to fully co-activated modules. We investigate how the temporal structure influences the network’s capacity to encode and selectively retrieve multiple dynamical patterns, while considering biological constraints such as the cost of long-range connectivity. Our results offer insight into how spatiotemporal coding and network organisation support robust, large-scale memory storage and replay.

## I. INTRODUCTION

Healthy brain function emerges from a rich dynamical repertoire, reflected in spontaneous and cue-evoked replay of spatiotemporal activity patterns spanning microscopic to large scales [1–7].

Memory replay - the reactivation of activity configurations that occurred during previous, active experience - was primarily observed during hippocampal sharp-wave ripples (SWRs) in sleep [8], but has also been reported during wakefulness and in cortical regions beyond the hippocampus [9–13]. Notably, also, scale-free neuronal avalanches have been shown to consist of repeating patterns of precise spiking [14], with single neurons participating selectively in specific LFP-based avalanche patterns [15]. Replay is thought not only to support memory consolidation but also to enable memory recall during task performance, providing the building blocks for mental simulation, planning, and prediction [16–18].

The mechanisms by which a given stimulus selectively engages specific subnetworks, leading to the reactivation of the appropriate learned sequences in response to a specific environmental stimulus, are not yet fully understood. Notably, stimulus-triggered reactivation of spatio-temporal patterns of activity, which reflect patterns observed during spontaneous activity, has been observed in vivo [4, 19–21], in vitro [19, 22, 23], and in in-silico [24].

Several results indicate that the contribution of replay to memory processes involves multiple spatial and temporal scales [25–28]. Further evidence on the crucial role of precisely coordinated spatiotemporal activity in neuronal assemblies comes from experiments on spike-phase coding in the auditory and visual primary cortices [29, 30] and from experiments on the short-term memory of multiple objects in the prefrontal cortices of monkeys [31]. While memory formation mechanisms, such as spike-timing-dependent plasticity, operate at the molecular and synaptic level, the execution of complex tasks requires the coordination of activity across regions at both mesoscopic and large-scale (whole-brain) levels [3]. However, long-range connections are costly, and the brain must operate within energetic constraints [32]. This creates a need to reach an optimal balance: maximising the number of retrievable, large-scale patterns while minimising the structural and energetic costs of inter-module communication.

Thus, we aim to investigate cue-evoked replay of multiscale spatiotemporal patterns and to determine the extent to which the simultaneous engagement of multiple regions by a process is necessary to achieve optimal multiscale coordination (i.e., maximise the number of stored patterns that can be selectively and successfully replayed) at the minimum cost (in terms of utilising large-scale connections). To do this, we build upon a previously designed modular neural network [24] composed of leaky integrate-and-fire (LIF) neurons, inspired by Hopfield-like associative memory systems for dynamical patterns [33] and governed by a spike-timing-dependent plasticity learning rule. Numerous experimental studies have shown that synaptic efficacy can be bidirectionally modified: it can be strengthened through long-term potentiation (LTP) or weakened through long-term depression (LTD), depending on the precise relative timing of pre- and postsynaptic spikes [34–37]. These seminal observations have been repeatedly confirmed and have motivated the development of phenomenological models of spike-timing-dependent plasticity (STDP) which describe how LTP and LTD depend on both the order and the temporal separation of pre- and postsynaptic action potentials [38–46].

The model is designed to learn and store a series of spatio-temporal patterns, defined at the mesoscopic level as ordered sequences of activated modules, and at the microscopic level as precise spike sequences across neurons. Learning is achieved through appropriate modifications of excitatory and inhibitory weights connecting the neurons. Starting from a set of disconnected neurons grouped into modules, the spatiotemporal paths progressively shape synaptic connections corresponding to neurons spiking in sequence, by means of the STDP learning rule. This way, the information is encoded both spatially, through the selection of active neurons as in the Hopfield model, and temporally, through the specific spiking order and phase relationships among active units. As the encoded patterns are organised as travelling waves among modules, the resulting connectivity matrix will reflect this modular organisation. We introduce a parameter, denoted *η*, named the “module co-activation index”, which controls the extent of temporal overlap between module activations and, consequently, the topology of the resulting connectivity matrix.

This parameter regulates the width of the distribution of spiking times of the neurons in a specific module, and therefore the strength of the oscillatory activity of the module in a given pattern. The network can operate across distinct temporal regimes depending on the value of *η*, ranging from a perfectly ordered sequential activation of modules to a regime where modules are active simultaneously, while still preserving precise spike timing at the level of individual neurons. In particular, when modules fire sequentially with zero or minimal temporal overlap, the network behaviour resembles a synfire chain. The synfire chain is characterised by a feedforward sequence of neuronal activations, where groups of neurons fire synchronously and propagate their activity to the next group, enabling the precise and reliable transmission of temporal information across circuits [47]. At high values of *η*, the temporal distinction between modules diminishes, leading to widespread co-activation and the loss of a clear sequential structure.

We investigate how varying degrees of the overlap between modules in the stored patterns affect the structural organisation of the resulting connectivity matrix and the network’s capacity to robustly store and retrieve patterns across a range of dynamic regimes, including cue-driven and spontaneous dynamics.

## II. THE MODULAR LIF MODEL

We investigate how the modular architecture of spiking neural networks influences their capacity to store and retrieve dynamic activity patterns, defined both at the microscopic level (precise spike timing) and the mesoscopic level (sequences of module activations). The network is composed of *S* modules, each containing *Z* leaky integrate-and-fire (LIF) neurons, yielding a total of *N* = *Z S* neurons. A set of *P* dynamic periodic patterns, travelling waves among modules with precise spatio-temporal spiking structure, is stored in the network during a learning stage. The network is then tested to assess whether a partial cue can selectively trigger the replay of the corresponding pattern.

In the learning stage, the synaptic strengths *J*_*ij*_ of connections between the pre-synaptic neuron *i* and post-synaptic neuron *j* are established according to the following STDP-based learning rule:

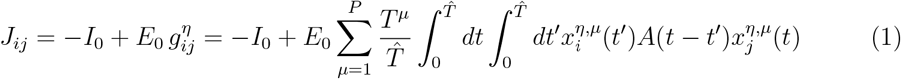

where *I*_0_ is a global inhibitory term, and 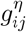 is the structured component induced by the learning of *P* dynamical patterns, whose modular organization is determined by *η*. The sum over *µ* runs over all *P* patterns to be stored,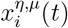 is the spiking activity of the *i*^th^ neuron in pattern *µ*, with the module co-activation indexed by *η. T*^*µ*^ is the period of the pattern *µ* to be stored, and the learning time 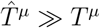 is much longer than the individual pattern period *T*^*µ*^, allowing for sufficient integration over multiple cycles. *E*_0_ is a parameter that determines the strength of the structured term, and the function *A*(*t*− *t*^*′*^) is the spike-timing dependent plasticity (STDP) learning window, which determines the weight modification when a time delay *τ* = *t* − *t*^*′*^ occurs between pre- and post-synaptic spikes, shown in Fig. 1, defined as:

**FIG. 1:**
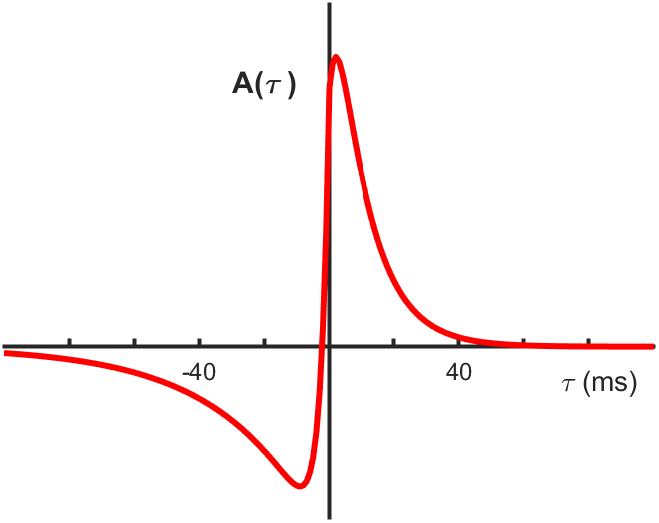
Learning window A(*τ*) used in the learning rule (1) to model STDP, showing potentiation for positive *τ* and depression for negative *τ*, fitted [42] to experimental data from Bi and Poo [35]

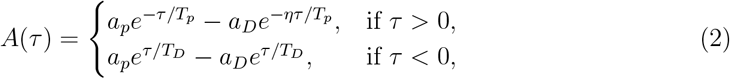

with *a*_*p*_ = *A*_0_*/*1 + *ηT*_*p*_*/T*_*D*_, *a*_*D*_ = *A*_0_*/η* + *T*_*p*_*/T*_*D*_. The function parameters are the one introduced in [42], to fit the experimental data described in [35], and used in [24, 33, 48, 49]. It is such that the connection between pre- and post-synaptic units is potentiated if the pre-synaptic neuron fires a few milliseconds after the post-synaptic neuron, and depressed if the order is reversed. The normalisation 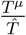 ensures that the connections do not depend on the learning time 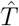, if it is large enough that the border effect can be neglected. Due to the biological constraint that neurons in real neural circuits are either excitatory or inhibitory, synapses arising from excitatory neurons must be positive, while those from inhibitory neurons must be negative. Starting from this, in our model, the effective synaptic coupling between neurons *i* and *j* is composed of two parts: (i) a negative global inhibitory contribution, produced by a fast inhibitory population (see the discussion on the origin of global inhibition in [50]), and (ii) a learning–dependent term *E*_0_*g*_*ij*_. During the initial learning procedure, P periodic spatiotemporal patterns are stored in the network connections. After the learning phase, the quantities *E*_0_*g*_*ij*_ associated with the same presynaptic neuron *j* may take either positive or negative values. However, *E*_0_*g*_*ij*_ should be interpreted as a modification of an initial baseline synaptic efficacy, which we assume to be identical for all pairs of neurons and denote by *J*_0_. The full synaptic weight is therefore written as (*J*_0_ + *E*_0_*g*_*ij*_). The constant *J*_0_ is chosen such that, after learning, all synapses arising from excitatory neurons remain strictly positive:

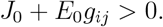

With this choice, the compensating inhibitory term becomes − (*I*_0_ + *J*_0_), and the expression in Eq. (1) can be rewritten as

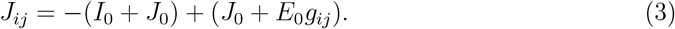

This formulation ensures that all synaptic connections between units *i* and *j* remain excitatory and positive, while a global inhibitory contribution maintains network balance. As a consequence, the system consists of *N* excitatory units with positive mutual couplings, and global inhibition preserves equilibrium [33].

The dynamics of the network are simulated considering a leaky integrate-and-fire model of neurons, where the membrane potential follows the equation:

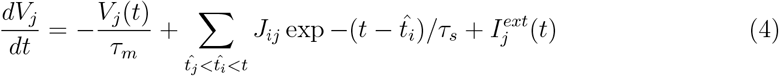

where sum on 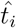 runs on all the spikes in input to the neuron *j* after the last spike of neuron *j*, 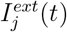 is the external input, *J*_*ij*_ is the result of learning process in eq.1, the membrane time constant is *τ*_*m*_ = 10*ms*, the synapse time constant is *τ*_*s*_ = 5*ms* [51, 52]. When the potential reaches the threshold Θ (conventionally set to Θ = 1), the neuron fires and the potential is reset to the resting value *V*_*j*_ = 0. Note that, among the parameters *I*_0_, *E*_0_ and Θ, only two are independent. We have chosen to set Θ = 1, but one could as well set *E*_0_ = 1, for example, and study the dependence of the model with respect to *I*_0_ and firing threshold Θ.

Each pattern 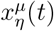, serving as a teaching signal for cortical plasticity, is a periodic sequence of spikes involving only *G* < *S* modules, and only *K* < *Z* neurons inside each active module. For each pattern, an ordered sequence of *G* active modules

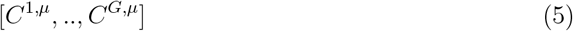

is randomly selected, while the remaining modules remain silent. In the following, the number of modules is set to *S* = 66 with *Z* = 200 neurons per module. The number of active neurons per pattern, *GK*, is fixed at *SZ/*4. The same module can participate in different patterns and appear multiple times within the same pattern. We consider two configurations satisfying this condition: (i) *G* = *S/*2 = 33 and *K* = *Z/*2 = 100, representing a wave spanning half of the modules; and (ii) *G* = *S* = 66 and *K* = *Z/*4 = 50, representing a wave spanning all modules. The choice of the parameters *G* and *K* is guided by empirical observations and theoretical considerations. Analysis of source-reconstructed resting-state MEG data from humans [24] shows that long-lasting neuronal avalanches typically involve about half of the cortical regions defined by the Desikan–Killiany–Tourville atlas, corresponding to approximately 33 out 66 active modules. Accordingly, we explore values of *G* in this range, including *G* = 66 as a limiting case of full-pattern propagation. The number of active neurons per module *K* was chosen so that each pattern recruits about 25% of the total network units, in line with theoretical results showing that maximal information capacity is achieved when only a fraction of neurons participates in each stored pattern [33]. The degree of temporal overlap between activations of different modules is controlled by the parameter *η*. To each active neuron in pattern *µ* is assigned a phase of spike 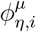 in the interval [0, 2*π*] that depends on index *η* and on the module to which neuron *i* belongs. Let’s denote by *C*_*i*_ the module to which neuron *i* belongs, and by *C*^*k,µ*^ the *k*-th module in the cyclic sequence defining pattern *µ*. Then the *P* spatiotemporal pattern 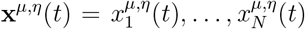, with *µ* = 1, …, *P*, are defined as:

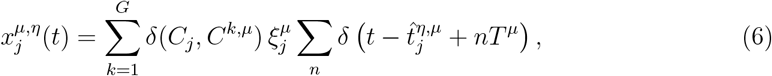

where:

*δ*(*C*_*j*_, *C*^*k,µ*^) is the Kronecker delta, equal to 1 if neuron *j* belongs to the *k*-th active module in pattern *µ*, and 0 otherwise;

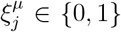 is set to 1 only for *K* randomly chosen neurons in active modules and 0 otherwise;

*T*^*µ*^ is the period of the stored pattern; we use *T*^*µ*^ = 125 ms ∀*µ*, corresponding to an 8 Hz alpha rhythm. Given the shape (i.e., temporal width) of the STDP kernel used in the model, a 125 ms period ensures that STDP remains sensitive to phase differences *within a single oscillatory cycle*. This is usefull for the type of dynamics we aim to capture: even relatively small variations in spike phase lead to distinct synaptic modifications, and the chosen period allows the model to reliably reflect these differences.

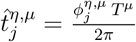 is the spike timing of neuron *j* in pattern *µ*, where 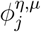 denotes the phase of firing of neuron *j*, which depends on its module *C*_*j*_ and on the module co-activation index *η* as detailed below.

Therefore, from eq.1 and 6, we get the microscopic structured connectivity matrix *J*_*ij*_ among the *N* = *ZM* neurons:

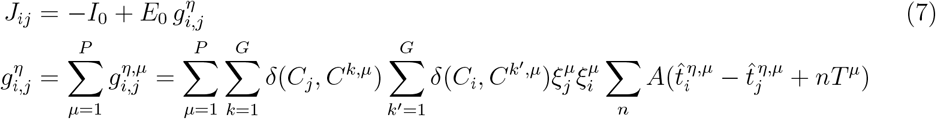

The phase of firing 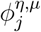, and therefore the spiking time 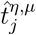, in pattern *µ* of neuron *j* belonging to module *C*_*j*_, is sampled from a probability distribution that allows tuning the extent of temporal overlap across modules in the wave through the index *η*. Specifically, *η* controls the extent to which spikes within each module are temporally spread across neighbouring modules. A “module distance” is defined as 2*π/G* in phase space, or equivalently *T*^*µ*^*/G* in the temporal domain, where *G* is the number of active modules in the pattern and *T*^*µ*^ its period. More precisely, for each module involved in a given pattern, the *K* spikes are drawn from a Gaussian distribution centred at the “centre phase” of the module. The standard deviation of this distribution is set to *η* times half a module, i.e.,

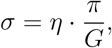

thus controlling the degree of temporal overlap between consecutive modules.

As an example, when *η* = 0, the spikes within each module are perfectly synchronous, and perfectly phase-shifted relative to the next module by 2*π/G*, forming a strictly sequential activation pattern, as in synfire chains. As *η* increases, spike times become increasingly dispersed around the centre of each module. For values of *η* large compared to *G/*2, the activation profiles of all *G* modules in eq.5 become very broad, resulting in flattened and overlapping co-activations across all the active modules. As a consequence, the original sequential structure of the pattern is no longer recognisable. This effect is clearly visible in Fig.2 panels (a,b), which shows an example of a stored pattern for four increasing values of *η*, and it also persists in the emergent dynamics during cue-induced retrieval of patterns as shown in Fig.4 (Fig. 1 of Supplementary Material shows the encoding of another modular spatio-temporal pattern during learning). Each panel in Fig.4a shows a raster plot of neuron spiking activity over time. At *η* = 0, neurons within each module are highly synchronised, firing almost simultaneously, and the activity of each module is well separated. As *η* increases, neurons belonging to a given module begin to fire before the activity of the preceding module has fully ceased, leading to increasing overlap between the activity of different modules. This behaviour is also evident in the firing rate of each module (panels in b). At low values of *η*, module activation is oscillatory and temporally well-separated, highlighting a clear sequential structure among modules. As *η* increases, module activities increasingly overlap, and their firing rates become less temporally distinct.

After the learning stage, the dynamics of the network started by a short cue are studied. A partial cue consisting of *H* spikes is presented, i.e., a short train of spikes is externally imposed on a small subset of neurons (*H* ≪ *N*), following the order *ϕ*_*h*_, with *h* = 1, …, *H*, as defined by one of the stored patterns 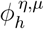, and using a period denoted by *T* ^cue^. In the following the period of the stored patterns is *T*^*µ*^ = 125 ms ∀*µ, H* = 75, and *T* ^cue^ = 83 ms. Specifically, the cue’s duration is less then or equal to (*HT* ^cue^)*/*(*GK*), and much less than one period.

The cue-induced emerging collective dynamics are analysed as a function of the number of stored patterns *P* and the module co-activation index *η*, to assess whether the network can reliably and persistently reproduce the pattern associated with the cue.

## III. RESULTS

We first characterize the connectivity structure produced by the STDP-inspired learning rule, as described in the previous section, and then the resulting recall dynamics, clarifying how the dynamic activity patterns, defined in terms of precise spike timing at the microscopic level and partially overlapping sequences of module activations at the mesoscopic level, are represented in the network. Thus, the dynamic activity patterns depends on the modular structure of the patterns, parameterised by *η*, which in turn shapes the modular organization of the learned connectivity.

The learning procedure in eq.7 generates a connectivity matrix that remains fixed during network dynamics. The number of patterns *P* and parameter *η* play a crucial role in shaping the network topology. As *η* increases, the network gradually changes from a strongly modular architecture to a more homogeneous structure.

The network connectivity at two values of *η* is shown in Fig. 3 for two different configurations (*G* = 33, *K* = 100) (top panels) and (*G* = 66, *K* = 50) (bottom panels).

To measure the degree of modularity of the microscopic connectivity 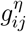 w.r.t. the partition into *M* modules, we check whether neurons within the same module exhibit stronger excitatory connections among themselves compared to their connections with neurons in other modules. We therefore define a *modularity measure*, following [24], as

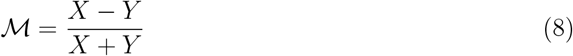

where *X* represents the average strength of internal positive connections within modules, while *Y* denotes the average strength of positive connections to neurons external to the module:

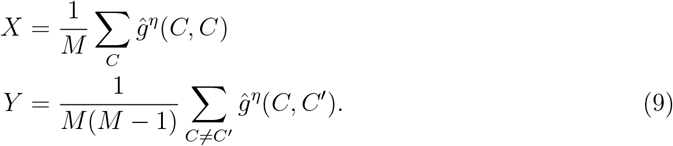

where the effective mesoscopic connectivity matrix between modules *ĝ*^*η*^(*C, C*^*′*^), is obtained summing excitatory connections over neurons belonging to modules *C* and *C*^*′*^, i.e.:

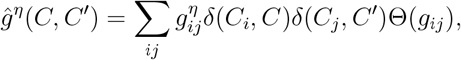

The value of ℳ can range in [− 1, 1]: it is positive if the strength of internal connections is greater than the expected value for a random network, and negative otherwise. In a random network, ℳ equals zero on average; conversely, in a network with completely disconnected modules, it equals 1. Here, with *P* = 100 patterns stored we observe ℳ _4_ = 0.8686 and ℳ _66_ = 0.1766 respectively for *η* = 4, 66, at *G* = 33, *K* = 100 and ℳ _4_ = 0.8314 and ℳ _66_ = −0.006 at *G* = 66, *K* = 50 shown in Fig. 3.

At low values of *η* (*η* = 4), the connectivity matrix *g*_*ij*_ exhibits a pronounced modular structure that reflects the tendency of neurons within the same module to fire in close temporal proximity, leading to strong intra-module connections through STDP. This modularity is also evident on the mesoscale, as seen in the coarse-grained excitatory matrix between modules *ĝ*^*η*^(*C, C*^*′*^) in Fig.3 panels (a), (c) on the right, which shows a clear diagonal structure indicative of strong intra-module connectivity. Positive connections also appear off the diagonal (even at *η* = 0 - see Fig. 2 of Supplementary Material), since the LTP window of STDP rule and the structure of patterns are such that connections between different modules are potentiated. Indeed, the time lag between spikes of consecutive modules (time lag *dt* = *T*^*µ*^*/G* at *η* = 0) lies well within the LTP window of our STDP rule (Fig.1). Conversely, if G is reduced and *T*^*µ*^ is increased excessively, the distance between spikes of consecutive modules may fall outside the STDP range, and the learning of the pattern would fail at low *η*. This does not occur at larger *η* since overlapping module activity brings spikes of different modules within the STDP window, allowing inter-module connections to be strengthened. As *η* increases (*η* = 66), the network becomes significantly more homogeneous. This is reflected in the diminished diagonal structure in both the full connectivity matrix and the coarse-grained one, signaling a near-complete loss of modularity and the emergence of widespread, long-range connections. Notably, the effect of high *η* differs depending on the network configuration. When only a subset of modules is involved in the pattern (panel (b), *G* << *S*), a residual modular structure persists: the modules participating in the pattern form intra-module connections between them, but not with modules not active in the pattern. In contrast, when all modules are active and *η* reaches the size of the network (panel (d)), the connectivity matrix becomes almost homogeneous on the scale of the whole network. In this case, synaptic weights are uniformly distributed across the network, and no modular organisation remains.

**FIG. 2:**
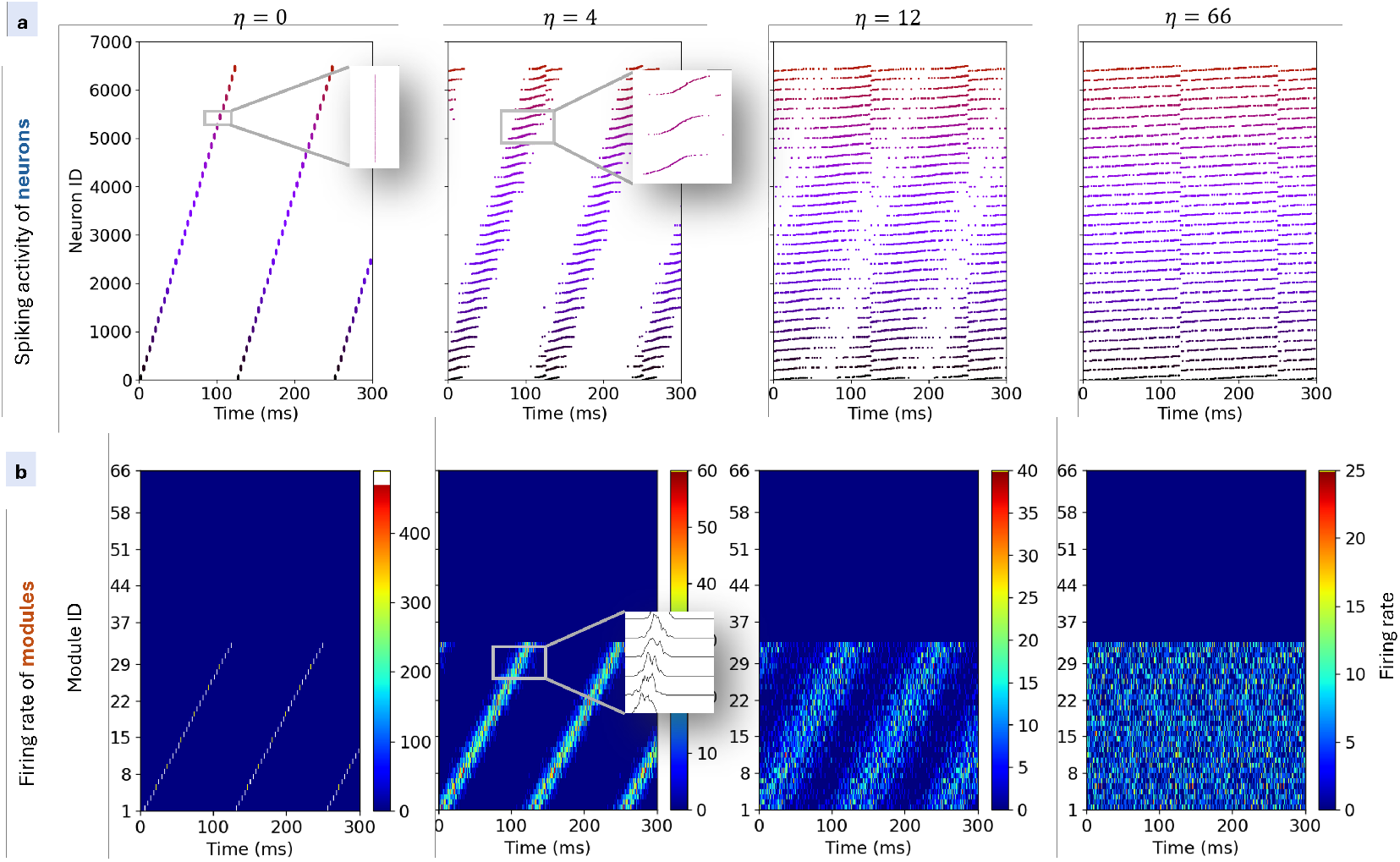
Encoding of modular spatio-temporal pattern during learning. Each stored periodic pattern is an ordered sequence of *G* ≤ *S* active modules, with overlap between modules regulated by *η*, and precise spike timing for the *K* ≤ *Z* non-silent neurons. Activity at increasing *η* values (0,4,12,66) is shown with *G* = 33, *K* = 100 for one specific pattern. Each panel in (a) shows a raster plot of the first 7000 neurons (while others are silent), with neurons sorted according to the pattern phase inside each module. At *η* = 0, neurons within each module are highly synchronised, firing almost simultaneously, while each module is engaged sequentially. As *η* increases, spike timing between groups becomes less distinct. In other words, neurons belonging to a given module begin to fire before the activity of the preceding module has entirely ceased, leading to increasing overlap between the activity of different modules. For high *η*, firing becomes broadly distributed with substantial temporal overlap between neuronal modules, reflecting a more homogeneous activation across the network. This behaviour is also visible in panels (b), showing the firing rate of each module, computed using 1 ms bins and expressed in Hz per neuron.

**FIG. 3:**
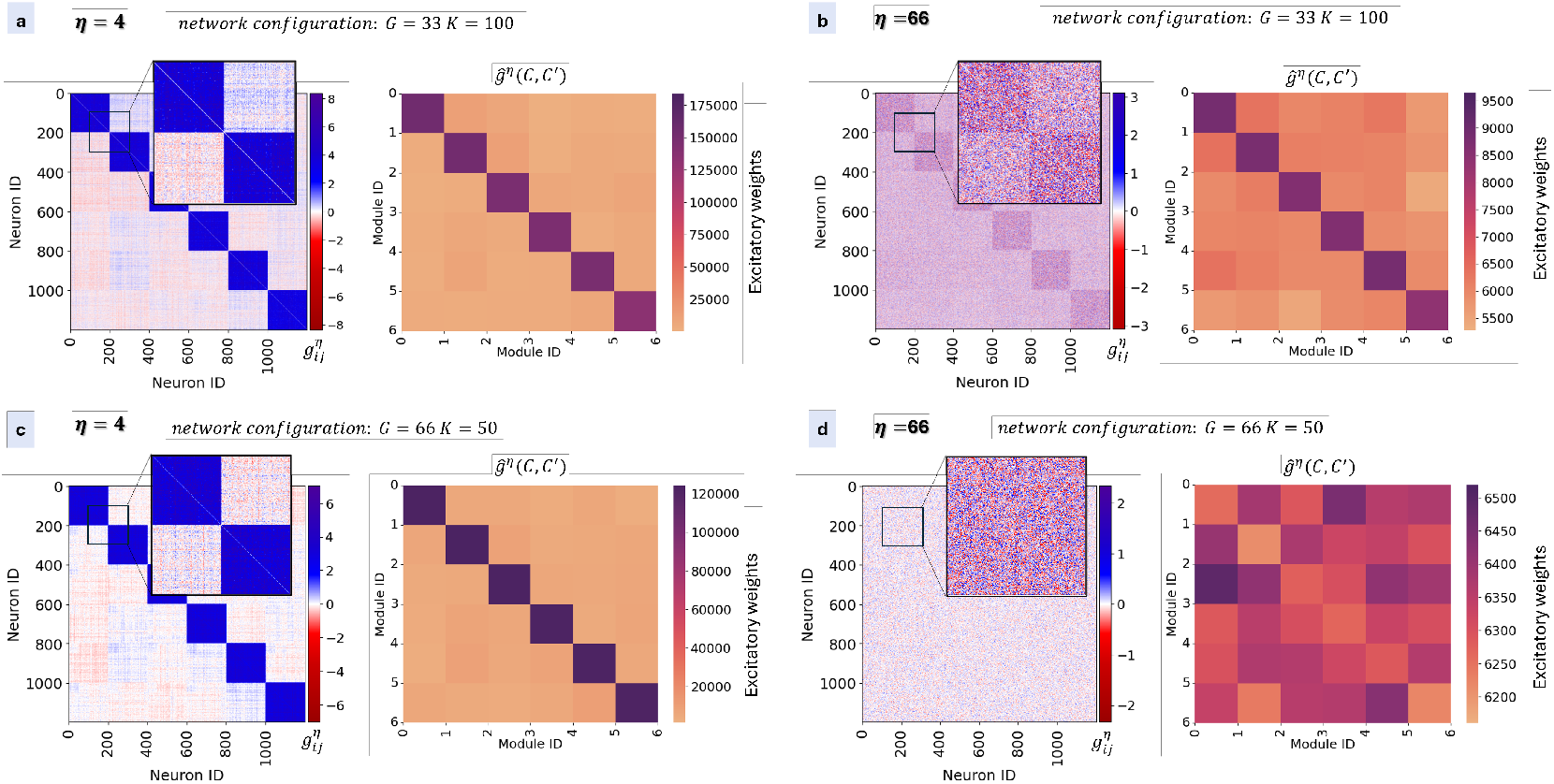
Effect of *η* on the network connectivity for two different configurations of the network. Panels (a) and (b) correspond to the network configuration with (*G* = 33, *K* = 100), shown for *η* = 4 and *η* = 66, respectively, and the network was forced to learn *P* = 400 patterns. Panels (c) and (d) display the configuration with (*G* = 66, *K* = 50) for the same values of *η*. Each panel shows the full connectivity matrix 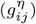 between individual neurons (left), with only the first 1200 neurons (corresponding to the first 6 modules) shown for readability, and the coarse-grained excitatory connectivity matrix (*ĝ*^*η*^(*C, C*^*′*^)) between modules (right), where only 6 of the 66 modules are displayed. At low *η* (*η* = 4), the network exhibits strong modular organisation: positive synaptic weights 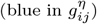 are concentrated within modules, reflecting the temporal clustering of neuronal activity due to STDP. This structure is preserved at the mesoscale in *ĝ*^*η*^(*C, C*^*′*^), where the diagonal dominance indicates strong intra-module connections. As *η* increases (*η* = 66), the connectivity becomes more homogeneous: synaptic weights are more evenly distributed across the network, and the diagonal structure in both 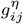 and *ĝ*^*η*^(*C, C*^*′*^) is reduced, indicating a loss of modularity and the emergence of long-range connections. In the (*G* = 33, *K* = 100) case, where only a subset of modules participates in the pattern (panel (b)), a residual modular structure persists even at high *η*, in opposition to when all modules participate in the pattern (panel (d)).

### Analysis of cue-induced dynamics

To investigate the storage capacity of the network as a function of the module co-activation index *η*, we examine the cue-induced emerging collective dynamics after applying a brief cue stimulus, to evaluate whether the network can selectively and persistently reproduce the pattern associated with the cue. To quantify the accuracy of these replays, following [33], we define an order parameter *q*^*µ*^ that measures the similarity between the emergent spiking activity and the stored pattern *µ* associated with the cue:

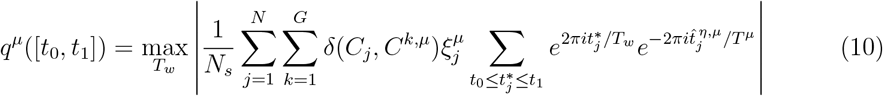

where the sum is taken over spikes *t*^∗^ emitted during a specific time window [*t*_0_, *t*_1_], 200 ms long, and factor 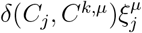 in the sum ensures that only neurons actively belonging to pattern *µ* contribute to the sum, while the “wrong firings” of other neurons affect the measure through the normalisation by the total number of emitted spikes *N*_*s*_. When only the correct neurons belonging to the correct modules fire, and there exists a replay window *T*_*w*_ for which their phases closely match the encoded ones, the exponential terms approach 1, causing *q*^*µ*^([*t*_0_, *t*_1_]) to approach 1. The “probe” time window *T*_*w*_ is the duration of the interval used to assess whether the collective dynamics match one of the stored patterns, as described in [53]. While the stored patterns are periodic with an intrinsic period *T*^*µ*^, during cue-induced dynamics, the network may replay them at compressed or dilated timescales. Therefore, the effective replay period is a priori unknown. The order parameter, defined as the maximum of the overlap across the tested windows *T*_*w*_, captures whether any segment of the spike train exhibits the correct phase ordering, effectively estimating the timescale of the replayed pattern. Therefore, measuring the order parameter also provides an estimate of the recall period 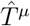.

Panels (a) and (b) of Fig. 4 examine the cue-evoked dynamics when the connectivity matrix is constructed with *η* = 0, *P* = 2 and *η* = 132, *P* = 4 (at *G* = 33 and *K* = 100). Panel (a) shows the response of the network to a brief initial cue in the case of *η* = 0. A short pulse of *H* = 75 spikes, with the order corresponding to the first encoded pattern, is delivered to the network, leading to a self-sustained spatiotemporal activity. This activity corresponds to the specific pattern associated with the cue, as confirmed by the middle figure of panel (a), where the overlap *q*^*µ*^ is plotted over time and *q*^*µ*=1^ (corresponding to the first stored pattern) reaches and maintains a maximum value, indicating successful retrieval. The figure to the right shows the average firing rate for each module, highlighting perfectly sequential activations. Panel (b) shows a similar example for *η* = 132. While retrieval is still perfect in terms of precise spiking times, as also indicated by the high order parameter *q*^*µ*=1^, we observe that the dynamics no longer follow a well-defined sequential activation of the modules; instead, module activations begin to overlap. This suggests that, although the information is preserved at the microscopic level, as indicated by the high order parameter, it is lost at the mesoscopic level, since it is no longer possible to delineate the sequence of regions involved in the pattern (see the firing rate for each region). In Fig. 3 of the Supplementary Material, the effect of noise on cue-evoked replay is investigated. Recall is initiated by providing a small cue (as in Fig. 4), while different noise intensities are applied throughout the simulations to assess the robustness of pattern retrieval. We find that low levels of noise do not impair replay, whereas excessive noise disrupts it.

**FIG. 4:**
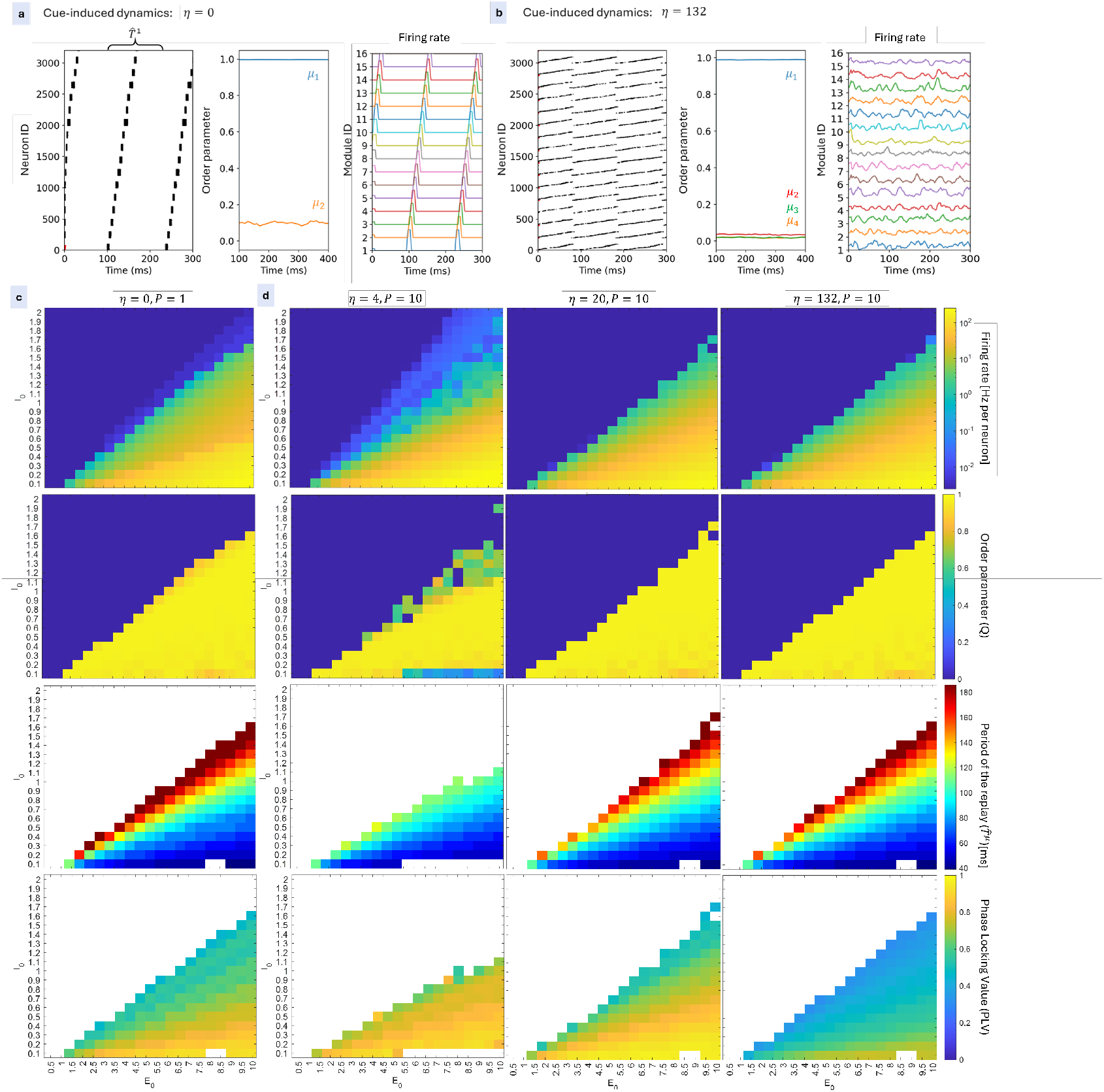
Cue-induced dynamics reveal the effect of *η* on pattern retrieval and spatiotemporal organization for the network composed by G=33 modules and K=100 neurons inside each module. Panels (a) and (b) show examples of cue-induced self-sustained spatiotemporal activity for networks whose connectivity matrices were constructed with (*η* = 0, *P* = 2) and (*η* = 132, *P* = 4), respectively. On the left of each panel, a raster plot displays spiking activity following a brief external cue (in red), composed of 75 spikes mimicking the first stored pattern (*µ* = 1). 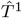 is the period of the replay of pattern 1. In the centre, the order parameter dynamics are shown: the overlap *q*^1^ reaches and maintains a value near 1, while the overlaps with other patterns remain low. On the right, the firing rate of each module computed using 1ms bins is shown. While retrieval is perfect in both cases, only for low *η* does the network preserve a clear sequential activation of modules. For high *η*, activations become increasingly overlapped, indicating a loss of structured dynamics at the mesoscopic level. Panels (c) and (d) show a systematic exploration of network emerging collective dynamics, after the cue presentation, as a function of global inhibition *I*_0_ and structured connection parameter *E*_0_. Panel (c) corresponds to the case with (*η* = 0, *P* = 1), while panel (d) displays results for *P* = 10 stored patterns with *η* = 4, 20, 132 (columns). Each row shows a different metric: (1) firing rate (expressed in Hz per neuron); (2) average order parameter *q*^*µ*^ to measure the quality of selective replay of cued pattern *µ*. The region of successful retrieval, at fixed P, becomes larger with *η*; (3) period of the replay (expressed in ms); and (4) phase locking value (PLV) computed as the average of the pairwise phase-locking values across all module pairs, which drops sharply as *η* increases, indicating reduced temporal coordination across modules despite successful microscopic pattern retrieval.

We then systematically repeat the cue stimulation using the first pattern (since all patterns are equivalent) across a range of network parameters to investigate how the ability to retrieve patterns depends on network excitability (*E*_0_) and global inhibition (*I*_0_), as well as on *η* (see eq. 12). Panel (c) shows the case of a single encoded pattern (*P* = 1) with *η* = 0, while panel (d) explores the case of *P* = 10 encoded patterns for *η* = 4, 20, and 132. At *P* = 10 the case *η* = 0 is not shown since there are no value of (*I*_0_, *E*_0_) with successfull retrieval at *η* = 0, *P* = 10,*G* = 33, *K* = 100. Panels (c) and (d) display four key metrics used to characterise the system’s behaviour, arranged by rows. In the first row, the network firing rate of the cue-induced collective dynamics is shown. At a low value of *E*_0_ and a large *I*_0_, the firing rate drops to zero because the cue fails to trigger any activity. During replay, neurons do not necessarily emit a single spike per period: depending on the excitation–inhibition balance, multiple spikes per neuron can occur within each replay cycle, so that the mean firing rate is not uniquely determined by the replay period (see Fig. 4 of Supplementary Material). The second row shows the average order parameter over time, and we observe that for small *P* or large *η*, a broad region in the parameter space (*E*_0_, *I*_0_) exhibits *q*^*µ*^ ≈ 1, indicating successfull retrieval. At fixed *P*, as *η* increases, this region expands and then stabilises, showing no small changes between the last two shown values of *η*. The third row displays the period of the replayed pattern. Notably, the period of the replayed pattern can be tuned by changing *E*_0_ and *I*_0_, and is in general different from the period of the patterns used in the learning procedure. Generally, it increases by increasing the global inhibition *I*_0_, or increasing *E*_0_. In the fourth row, to better characterise the consistency of the temporal coordination between different regions of the network, we compute the Phase Locking Value (PLV) [54], which captures the differences in spatial organisation of the recalled dynamics. To this end, we computed the firing rate of the modules in 10*ms* time bins. The pairwise phase relation between the activity of two given modules *C* and *C*^*′*^ is defined as:

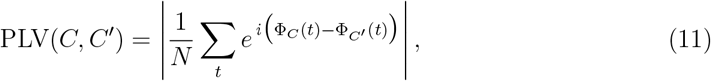

where Φ_*C*_(*t*) is the instantaneous phase extracted using the Hilbert transform [55] of z-scored rate signal of the module *C*. The average over all module pairs is then computed and denoted as the PLV. As *η* increases, despite the high order parameter values, the PLV drops significantly, indicating degraded phase-locked synchronisation across modules. This suggests that, although the information is preserved at the microscopic level, it is lost at the mesoscopic level, where the spatial sequence of activation can no longer be clearly delineated.

### The storage capacity of the network

Here, we investigate how the structural organisation, shaped by *η*, translates into the ability to store and selectively retrieve many spatiotemporal patterns, as opposed to the previous section, where the behaviour of the system was addressed for a fixed number of stored patterns. By systematically stimulating the network with brief cues, we evaluate the network’s storage capacity, denoted as *P*_max_, for both network configurations, i.e. (*G* = 33, *K* = 100) and (*G* = 66, *K* = 50), as shown in Fig. 5. *P*_max_ is the maximum number of patterns that can be stored and successfully retrieved for a given value of *η*. In particular, we investigate the network’s storage capacity when a stimulus of *H* = 75 spikes, arranged in the order corresponding to pattern *µ* and with period *T*_*cue*_, is applied. Successful retrieval is defined as an overlap *q*^*µ*^ above 0.95, averaged over the last 400 ms of the simulation. It is evaluated for different values of *η*, numbers of stored patterns *P*, and parameters (*E*_0_, *I*_0_).

**FIG. 5:**
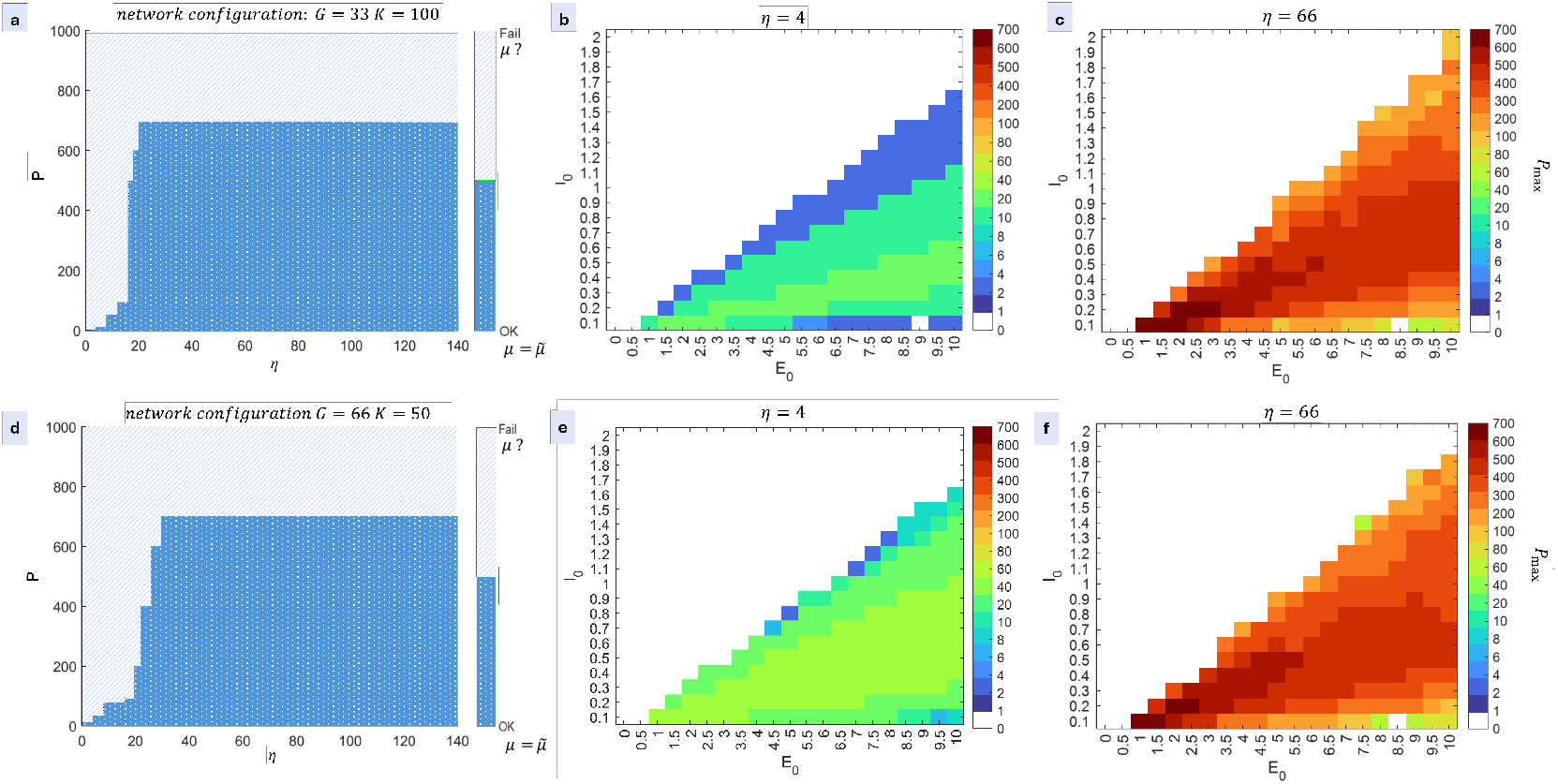
Effect of the module co-activation index *η* on the storage capacity of the network for two configurations. Panels (a–c) refer to the configuration (*G* = 33, *K* = 100), and panels (d–f) to (*G* = 66, *K* = 50). After storing *P* patterns, cue-induced dynamics is tested using a train of *H* = 75 spikes arranged according to pattern *µ* with period *T*_cue_. Successful retrieval is defined as an overlap *q*^*µ*^ > 0.95 with the cued pattern. Panel (a) shows the region in the (*P, η*) parameter space where there exists at least one combination of (*E*_0_, *I*_0_) for which the network can successfully and selectively replay the patterns (dark-blue region). Panels (b) and (c) show the maximum number of patterns *P*_max_ as a function of the parameters (*E*_0_, *I*_0_) at *η* = 4 and *η* = 66, respectively. Successful replay occurs in a parameter region that shrinks with increasing *P* when compared at fixed *η*, while, in general, higher values of *P* are attained at large *η*. Panels (d–f) show the same analysis for the (*G* = 66, *K* = 50) configuration.

Panel (a), corresponding to the case (*G* = 33, *K* = 100), shows the maximum capacity over all the values of (*E*_0_, *I*_0_), and for each given value of *η*. Two distinct regions emerge. A large dark-blue area, which expands with increasing *η*, corresponding to the zone where the network successfully works as an associative memory. In this regime, a cue triggers the correct replay of the corresponding pattern. A light-blue region, where the order parameter is close to zero, indicates that the network fails to produce activities related to the cued pattern. We observe that the successful replay region reaches the maximum *P* around *η* ∼ 20, with *P* = 700 stored patterns. The storage capacity is smallest at *η* = 0. When *η* = 0, information about the temporal order of neuronal spikes within each module is lost, and only the identity of the active neurons and the order of module activations are preserved. This loss of intra-module temporal structure increases interference between patterns that engage the same module, thereby reducing storage capacity.

In panels (b) and (c) of Fig. 5, we focus on two specific values of *η* to explore how the maximum number of perfectly recalled patterns depends on global inhibition *I*_0_ and the structured term strength *E*_0_. In panel (b) (*η* = 4), the maximum number of patterns that can be stored and successfully retrieved is limited to about *P* = 20. However, this value is not uniform across the (*E*_0_, *I*_0_) parameter space: the region where *P* = 20 patterns can be successfully retrieved is narrow, while large areas correspond to lower capacities. In panel (c) (*η* = 66), consistent with earlier results, the maximum storable number of patterns increases to about *P* = 700, achieved within a limited region of the parameter space. Nonetheless, lower numbers of stored patterns correspond to larger regions of (*E*_0_, *I*_0_). Panels (d–f) repeat the same analysis for the network configuration (*G* = 66, *K* = 50). The storage capacity increases with *η* as in the previous case and then stabilises; however, the plateau of the maximum number of patterns that can be stored and correctly recalled is reached at a higher *η* value. Panels (e) and (f) illustrate the maximum number of retrievable patterns as a function of (*E*_0_, *I*_0_) for *η* = 4 and *η* = 66, respectively. As in the case (*G* = 33, *K* = 100), for (*G* = 66, *K* = 50) successful replay also occurs in a parameter region that broadens with increasing *η* when compared at fixed *P*, and, overall, higher values of *P* are achievable at *η* = 66 than at *η* = 4. In Fig. 5 of Supplementary Material, the maximum number of patterns *P*_max_ is shown as a function of (*E*_0_, *I*_0_) at *η* = 20 for (*G* = 33, *K* = 100) and at*η* = 28 for (*G* = 66, *K* = 50), corresponding to the smallest *η* at which each network reaches the plateau of its maximum storage capacity.

### Trade-off between storage capacity and modularity of the network

Figure 6 illustrates the link between the topological reconfiguration of the network as a function of the parameter *η* and its ability to work as an associative memory. The first row characterises the network for the configuration (*G* = 33, *K* = 100). Panel (a,b) shows the average strength *X* of internal positive connections within modules, and the average strength *Y* of positive connections to neurons external to the module (see eq. (9)) as a function of *η*. As *η* increases, intra-module weight *X* decreases, especially when the network is initialised with a larger number of patterns (each line colour corresponds to a different number of stored patterns, *P*). At the same time, the average strength *Y* of long-range (inter-module) connections increases, which in turn affects the network topology. For example, the mean path length *L*, computed using Dijkstra’s algorithm to obtain the distance matrix from the weighted network structure, tends to decrease for high values of *η*, indicating that the network becomes more interconnected across modules. It is worth noting that in panel (c), curves for low values of *P* are missing, as the corresponding networks are too sparse to compute meaningful path lengths. To further explore the relationship, the modularity ℳ, defined in eq. (8), is computed and shown in panel (d) as a function of *η*. Higher values of ℳ indicate a stronger community structure, while values near zero reflect a loss of modular organisation. Modularity reaches its maximum at *η* = 0 and gradually decreases as *η* increases, eventually stabilising at a low asymptotic value. For the configuration (*G* = 33, *K* = 100), the point at which modularity becomes stationary is around *η* ≈ 30, which is close to the value at which the network’s memory capacity also plateaus, as observed in earlier analyses. This suggests a close relationship between structural modularity and functional memory performance. Notably, even for high values of *η*, the network retains a residual modular structure at (*G* = 33, *K* = 100) when only a subset of modules is involved in the pattern: STDP strengthens connections only between modules that are systematically co-activated within the same patterns, while modules not involved in those patterns do not develop strong inter-module connections, even for large values of *η*. In contrast, when all modules participate in the patterns ((*G* = 66, *K* = 50) in panel (g)), modularity approaches zero at very large *η*. This occurs because, when only a subset of modules is involved in the stored patterns, a distinction with silent modules remains, allowing the active modules to form well-defined communities with denser intra-community than inter-community connections. In both cases, the observed decrease of *M* with increasing *η* indicates a redistribution of synaptic resources that favours global integration over local cohesion.

**FIG. 6:**
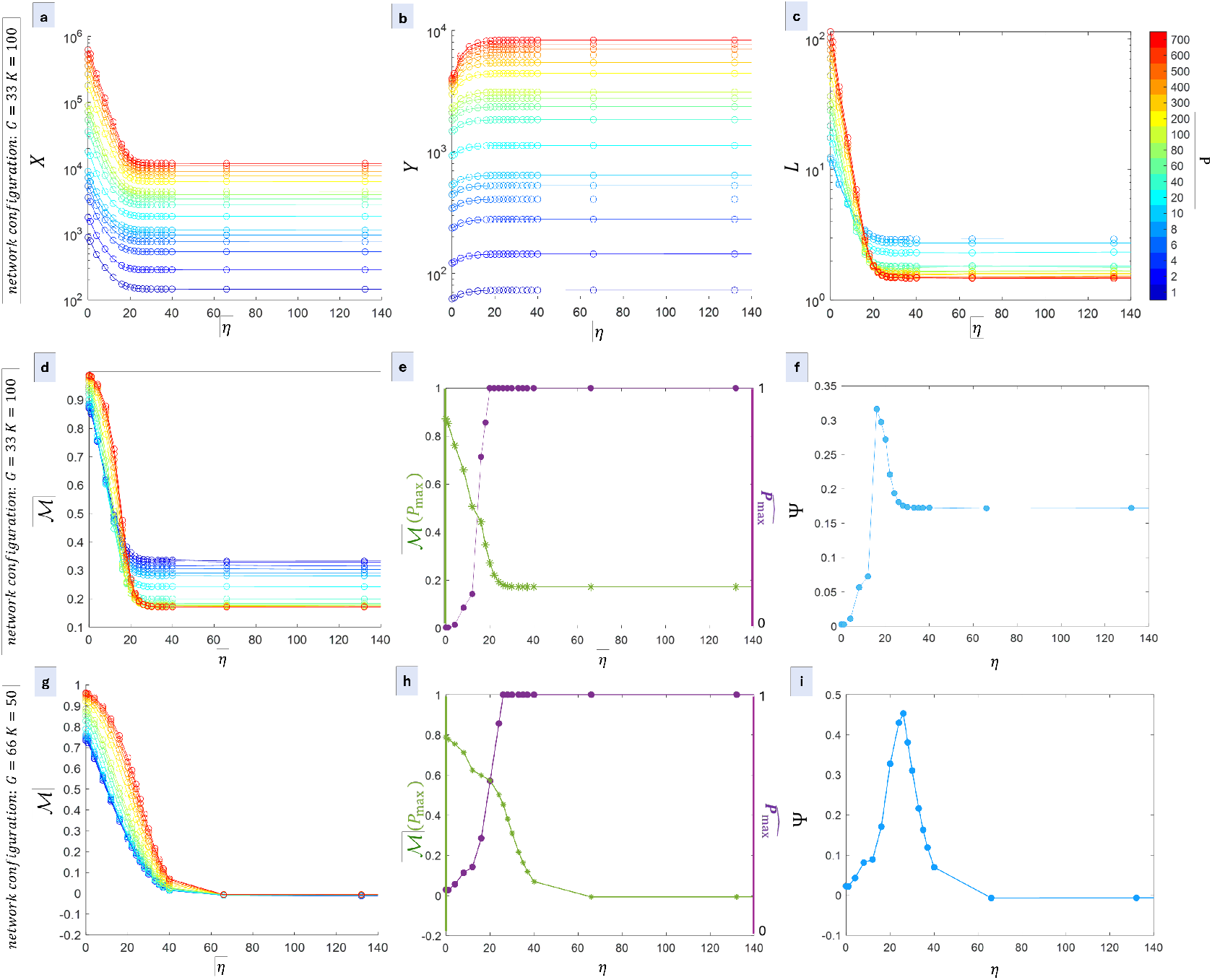
Role of module co-activation index *η* in structure and function: trade-off between wiring cost and storage capacity. Panels (a - f) show results for network configuration (*G* = 33, *K* = 100). Panel (a) displays the total strength of intra-module positive connections (*X*), which decreases with increasing *η*, especially for larger numbers of stored patterns *P* . Each colour line refers to a particular number of patterns used to build the connectivity matrix. Panel (b) shows the corresponding increase in inter-module (long-range) connection strength (*Y*), while panel (c) illustrates how the mean path length *L* decreases with *η*. Panel (d) reports the modularity ℳ, computed as in eq.8, which declines with increasing *η*, reaching a plateau around *η* ≈ 30. Panel (g) shows the same analysis for the network configuration (*G* = 66, *K* = 50). When all modules are engaged in the pattern, modularity approaches zero at high *η*, indicating the loss of excess intra-module connectivity relative to inter-module connections. Panels (e) and (h), corresponding to network configurations (*G* = 33, *K* = 100) and (*G* = 66, *K* = 50) respectively, show that normalised capacity (purple) increases with *η*, and the concurrent decline in modularity (green). Since long-range connections are costly, to quantify this trade-off between wiring cost and storage capacity, panels (f) report a fitness index Ψ, defined as the product between capacity and modularity, which peaks at intermediate *η* values. Panel (i), which refers to the (*G* = 66, *K* = 50) configuration, shows a similar trend. However, the peak of the fitness curve occurs at a slightly higher value of *η*, consistent with the fact that the storage capacity in this configuration also reaches its maximum at a higher *η* compared to the first network configuration.

Crucially, this reorganisation entails computational costs: long-range connections require the establishment of links between spatially distant units, which are energetically and structurally expensive. In the brain, such connections are sparse. Their sparsity underscores their selective importance in supporting efficient inter-areal communication while minimising wiring cost. Panels (e) and (h), corresponding to network configurations (*G* = 33, *K* = 100) and (*G* = 66, *K* = 50), respectively, show an increase in the normalised capacity (purple curve) with *η*, accompanied by a decline in modularity (green curve). The trade-off between wiring cost and storage capacity is captured by a fitness index Ψ, obtained as the product of capacity and modularity, shown in panels (f) and (i). This index peaks at intermediate values of *η*, indicating the existence of an optimal regime that preserves some modular structure while maintaining high memory performance. This finding suggests an optimal point with some degree of long-range connectivity sufficient to support global integration, yet limited enough to maintain the underlying modular architecture.

This optimum corresponds to the regime in which the structural economy of modular organization is best balanced with the functional requirement of retrieving multiple patterns. Notably, although high modularity reduces global retrieval capacity, modular structure may still confer a significant functional advantage that is not captured by *P*_*max*_. Indeed, the identity of the retrieved pattern can be reliably decoded at the module level, without relying on precise spike timing of individual neurons, when *η* is not too large, and therefore modularity is not lost. At intermediate values of *η* indeed, the population activity within each module remains sufficiently coherent so that the identity of the active pattern is visible in the modules’ activation profile. This functional advantage of modularity is not visible in Fig. 6, which focuses on the maximum number of attractor retrievals, but it becomes evident when examining module-level activity and phase-locking values (see Fig. 4).

### Analysis of spontaneous dynamics

To further characterize the network, we study spontaneous dynamics in the absence of cue stimulation, setting the input 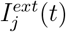 in Eq. (4) equal to 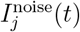, a noise term modelling spontaneous neurotransmitter release and other stochastic sources, defined as:

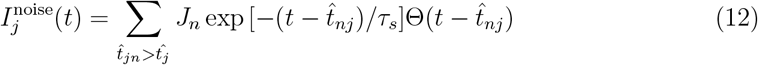

where *J*_*n*_ are random strengths extracted from a Gaussian distribution with mean 0 and standard deviation *α* = 0.5, independently for each neuron *j, τ*_*s*_ is the synapse time constant equal to 5 ms as in the Eq.4. The noise-event times 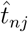 are also independently sampled for each neuron. The inter-event intervals *δt* follow a Poisson distribution:

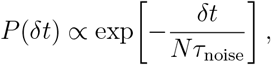

where *N* is the number of neurons and *τ*_noise_ is the characteristic timescale of the spontaneous activity. The rate of the noise per neuron, defined as *ρ* = 1*/*(*Nτ*_noise_), is equal to 1Hz [56].

Figure 7 shows the characterisation of spontaneous dynamics of the network for the first network configuration (*G* = 33, *K* = 100). Each row corresponds to a different value of *η* (*η* = 4, 20, 66), with *P* = 10 stored patterns. Each column refers to a different dynamic feature used to characterise the network’s spontaneous behaviour.

**FIG. 7:**
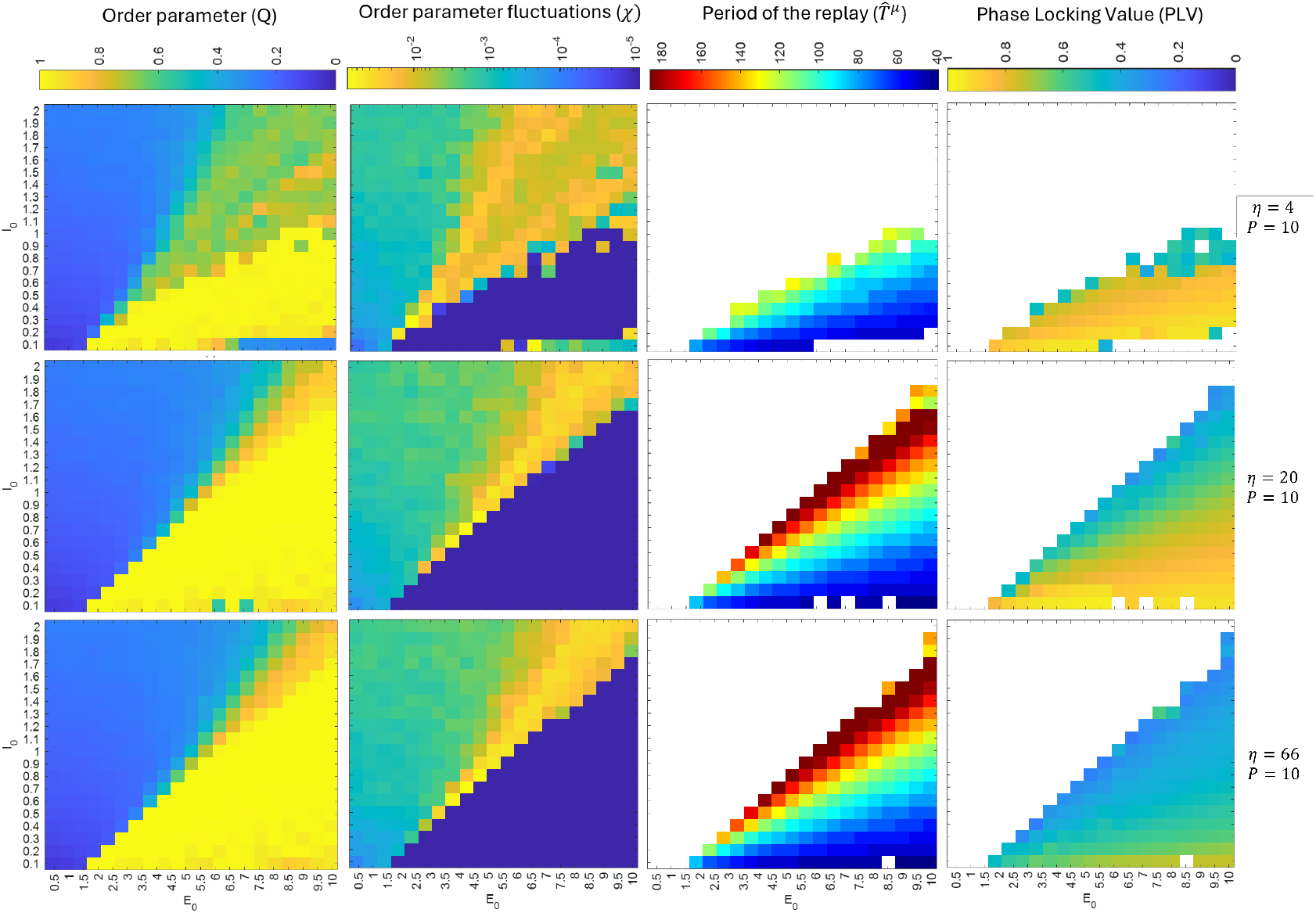
Spontaneous activity dynamics in the presence of noise for a network with (*G* = 33, *K* = 100) and *P* = 10 stored patterns. Each row corresponds to a different value of *η* (*η* = 4, 20, 66), and each column displays a different dynamic measure: order parameter, order parameter fluctuations, recall period, and phase locking value. The order parameter quantifies the similarity between spontaneous activity and any stored pattern, revealing regions where attractor states are spontaneously visited. At the edge between the region of successful spontaneous replay and the region with no-replay, there’s a region with large order parameter fluctuations, indicating a critical regime where alternation between high and low order parameter occurs in time. The recall period increases with *η*, reflecting the emergence ofattractors with varied frequencies of oscillation as a function of *E*_0_, *I*_0_. Phase locking decreases with *η*, showing reduced phase-locked synchrony across modules as long-range connectivity increases.

Here, in the absence of a cue, the order parameter is defined as the maximum of overlaps *q*^*µ*^ between the ongoing spontaneous activity and any of the *P* stored patterns, *Q* = max_*µ*_ *q*^*µ*^. Order parameter and its fluctuations *χ*_*Q*_, defined as the variance of the order parameter over time, are shown in the first and second columns, at different values of *η*. Consistent with previous findings [24], at the edge of the region exhibiting spontaneous self-sustained replay (identified by high values of the order parameter), a region emerges with large fluctuations in the order parameter. Order-parameter fluctuations provide a macroscopic indicator of the network’s propensity to transition between stored patterns, with larger fluctuations corresponding to increased dynamical flexibility, i.e. a greater number of distinct, transiently reactivated patterns, as described in [24] . A peak of the fluctuations, suggesting a critical regime [24], is clearly visible at intermediate and large *η*. Here, consistent with previous results [24], the network transiently and spontaneously alternates short replays over time, with spontaneous up/down fluctuations[57]. This is in line with the well-supported hypothesis, grounded in both experimental evidence and computational modeling, that the resting cortex operates in an extended dynamical regime, poised at the edge of a dynamical phase transition and exhibiting critical behaviour [58–65] The third column shows the period of the periodic pattern during the successful self-sustained replay. For low values of *η*, spontaneous replay is limited to a small region of the (*E*_0_, *I*_0_) space. In contrast, for higher *η*, the system spontaneously replays one of the stored patterns over a larger range of parameters, leading to a broader range of recall periods that vary as a function of the global inhibitory parameter *I*_0_ and the structural parameter *E*_0_. As in the cue-induced dynamics, increasing *η* leads to a decrease in phase locking activity of modules, suggesting that higher long-range connectivity allows for more distributed activity across modules, and a less precise sequence of module activations.

## IV. DISCUSSION

Many studies have investigated the role of modularity on collective dynamics [24, 66–69] and a substantial body of work has explored how STDP can be used to store and retrieve dynamical collective patterns [38, 43, 70–83]. In our work, we extend these lines of research by examining the interplay between modularity and replay, introducing a parameter that controls the degree of functional modularity of patterns.

This work builds on the framework introduced in [24], which explores a recurrent spiking neural network trained via spike-timing–dependent plasticity to store and spontaneously replay multiple spatio-temporal patterns. The network adopts a modular architecture, with each pattern engaging only a specific subset of modules. Modules represent mesoscopic units that can be interpreted anatomically as local neuronal populations (e.g., cortical regions or subregions) and functionally as units that play a specific role within a stored sequence. Extending this model, we incorporate the *module co-activation index η*, as a parameter controlling the degree of temporal overlap between module activities. The patterns presented as inputs during the learning stage encode information with precise temporal relationships at both the micro- and mesoscale level. Information is represented jointly by firing phase and firing rate, reproducing the structured and correlated population code observed experimentally[84–86]. These patterns consist of segregated modular cyclic spatio-temporal sequences (with the degree of segregation quantified by *η*), mimicking, for example, the segregated projections of different features into early sensory layers, where neurons in different modules exhibit distinct but partially overlapping tuning curves [87]. A sparse pattern-specific recruitment of neuronal subsets within each active module is assumed as a teaching signal during the learning stage, allowing us to focus on the consequences of such representations for learning, storage, and retrieval dynamics. Importantly, the functional modularity of the stored patterns directly shapes the structural modularity of the network connectivity emerging after the learning stage. Hence, *η* is meant to modulates the ability of regions to interact over the long-range, beyond local activities. These results talk to the converging evidence showing that computations might be conceptualized as either “local”, occurring primarily within regions, or “global”, where genuine interactions occur over the long range (e.g., utilizing long-range white matter projections), with mesoscale topological properties reflecting the balance between segregation and integration [88–91].

We analyse the effect of the module co-activation degree on the network storage capacity, defined as the maximum number of dynamical patterns that can be stored and selectively retrieved when a short cue is presented to the network. We show that the effects of temporal overlap extend beyond the network’s structural organisation, influencing its functional dynamics and storage capacity.

The concept of temporally overlapping activity reflects the biological nature of neural communication, which is distributed and temporally dispersed rather than perfectly synchronised. Neurons communicate through spikes that are distributed in time, and learning mechanisms such as STDP are sensitive to the precise spiking times. Changes in *η* not only modify structural properties, such as synaptic weight density, modularity, and path structure, but also shape the dynamics evoked by external cues, thereby affecting storage capacity.

For small *η* values, modules activate in a temporally distinct and sequential manner, enabling the detection of a sequential order across modules, and closely resembling the feedforward “synfire chain” architecture [47]. The synfire chains, characterised by their precise temporal synchrony, have traditionally been considered a biologically plausible model for sequence generation [47]. They are especially relevant in systems requiring temporally locked activity, such as the promotor HVC nucleus in songbirds, where neuron groups fire in tight sequences to support song production [92]. As *η* increases, temporal overlap between module activities grows, enhancing the network’s ability to store and retrieve a greater number of distinct spatiotemporal patterns, reaching a plateau. At very large *η*, the model approaches the one whose storage capacity was investigated in [33].

However, when *η* becomes larger, the sequential structure of patterns at the mesoscale is lost: module activations blur together, hindering the ability of the readout structure to decode pattern identity at the mesoscale level, namely through the temporal order of module activations. Moreover, maximising the number of retrieved patterns by increasing *η* produces a markedly less modular topology, reflecting the formation of numerous long-range inter-module connections. In a biologically realistic scenario, however, such connections are energetically costly, and the brain must balance memory capacity with metabolic constraints [32].

To quantify this trade-off, we track modularity, which inversely reflects the abundance of long-range connections, and define a fitness index Ψ as the product of capacity and modularity. Ψ peaks at intermediate *η*, identifying a regime where long-range connectivity is sufficient to support integration while preserving modular architecture.

Thence, the less modular neural network, emerging at intermediate *η*, exhibited a significantly greater capacity to store and recall multiple patterns, compared to the case *η* = 0, while still allowing the patterns to be decoded at the mesoscale based on the order of module activations—a possibility that is lost at very high values of *η*. This architecture, characterised by more distributed and long-range connections (with proportionally fewer intra-module connections), with respect to *η* = 0, can accommodate a broader variety of dynamical large-scale patterns. These findings are consistent with theoretical predictions from associative memory models and attractor networks, which emphasise the advantages of distributed representations and recurrent connectivity for memory capacity and robustness [93, 94]. Unlike the predominantly feedforward structure modelled by synfire chain networks, indeed the cortex appears to support more flexible, associative forms of memory that rely on recurrent interconnectivity and distributed representations [94]. Such properties are essential for higher cognitive functions, including learning, abstraction, and inference.

In conclusion, by introducing the fitness index that combines capacity and modularity, we identify an intermediate, “sweet spot” in which the network achieves both high memory performance and an economical topology. This balance between integration and segregation mirrors a principle often observed in cortical organisation, where the brain must store a rich repertoire of experiences while respecting energetic constraints. Our results suggest that effective associative memory systems, whether biological or artificial, can depend on operating in such a finely tuned regime, enabling the flexible, robust, and context-dependent retrieval of complex patterns that underpin higher cognitive functions. Hence, our findings show that memory performance in recurrent spiking networks is maximised at an intermediate degree of module co-activation. This regime reflects a fundamental principle of cortical organisation, enabling the brain to store a rich repertoire of experiences while remaining energetically efficient and flexibly adaptive.

## Author contributions

M.A., A.D.C., P.S., L.C., C.C, S.S. Conceptualization and Methodology; M.A., S.S., A.D.C. Simulations and Data analysis; M.A., S.S., P.S., C.C, L.C. Formal analysis; M.A., A.D.C., P.S., L.C., C.C, S.S. Writing - Original Draft; M.A., A.D.C., P.S., S.F., L.C., C.C, S.S. Writing - Review - Editing; M.A., A.D.C., P.S., S.F., L.C., C.C, S.S. Visualization; A.D.C., P.S., L.C., C.C, S.S. Supervision and Project administration; S.S., P.S., S.F. Funding acquisition; All authors read and approved the final manuscript.

## Acknowledgements

M.A., S.F., C.C., A.L., L.C. wish to acknowledge the Italian National Group for Mathematical Physics, GNFM-INdAM. S.F. wishes to acknowledge ICRANet.

## References

[1] K. Kaefer, F. Stella, B. L. McNaughton, and F. P. Battaglia, Replay, the default mode network and the cascaded memory systems model, Nature Reviews Neuroscience 23, 628 (2022).

[2] G. Deco, M. L. Kringelbach, V. K. Jirsa, and P. Ritter, The dynamics of resting fluctuations in the brain: metastability and its dynamical cortical core, Scientific reports 7, 3095 (2017).

[3] G. Deco, V. K. Jirsa, and A. R. McIntosh, Emerging concepts for the dynamical organization of resting-state activity in the brain, Nature reviews neuroscience 12, 43 (2011).

[4] R. Yuste, R. Cossart, and E. Yaksi, Neuronal ensembles: Building blocks of neural circuits, Neuron 112, 875 (2024).

[5] J. Cabral, M. L. Kringelbach, and G. Deco, Exploring the network dynamics underlying brain activity during rest, Progress in Neurobiology 114, 102 (2014).

[6] Y. Yang, D. A. Leopold, J. H. Duyn, and X. Liu, Hippocampal replay sequence governed by spontaneous brain-wide dynamics, PNAS Nexus 3, pgae078 (2024).

[7] S. Xu, W. Jiang, M.-m. Poo, and Y. Dan, Activity recall in a visual cortical ensemble, Nature Neuroscience 15, 449 (2012).

[8] M. A. Wilson and B. L. McNaughton, Reactivation of hippocampal ensemble memories during sleep, Science 265, 676 (1994).

[9] D. Ji and M. A. Wilson, Coordinated memory replay in the visual cortex and hippocampus during sleep, Nature Neuroscience 10, 100 (2007).

[10] D. R. Euston, M. Tatsuno, and B. L. McNaughton, Fast-forward playback of recent memory sequences in prefrontal cortex during sleep, science 318, 1147 (2007).

[11] K. Diba and G. Buzsáki, Forward and reverse hippocampal place-cell sequences during ripples, Nature Neuroscience 10, 1241 (2007).

[12] G. Girardeau and M. Zugaro, Hippocampal ripples and memory consolidation, Current Opinion in Neurobiology 21, 452 (2011).

[13] M. F. Carr, S. P. Jadhav, and L. M. Frank, Hippocampal replay in the awake state: a potential substrate for memory consolidation and retrieval, Nature Neuroscience 14, 147 (2011).

[14] T. L. Ribeiro, S. Ribeiro, and M. Copelli, Repertoires of spike avalanches are modulated by behavior and novelty, Frontiers in Neural Circuits 10, 16 (2016).

[15] T. Bellay, W. L. Shew, S. Yu, J. J. Falco-Walter, and D. Plenz, Selective participation of single cortical neurons in neuronal avalanches, Frontiers in Neural Circuits 14, 620052 (2021).

[16] A. Dimakou, G. Pezzulo, A. Zangrossi, and M. Corbetta, The predictive nature of spontaneous brain activity across scales and species, Neuron 113, 1310 (2025).

[17] H. F. Ólafsdóttir, D. Bush, and C. Barry, The role of hippocampal replay in memory and planning, Current Biology 28, R37 (2018).

[18] B. Liu and D. V. Buonomano, Ex vivo cortical circuits learn to predict and spontaneously replay temporal patterns, Nature Communications 16, 3179 (2025).

[19] A. Luczak, P. Barthó, and K. D. Harris, Spontaneous events outline the realm of possible sensory responses in neocortical populations, Neuron 62, 413 (2009).

[20] A. Luczak and J. N. MacLean, Default activity patterns at the neocortical microcircuit level, Frontiers in Integrative Neuroscience 6, 30 (2012).

[21] T. Livne, D. Kim, N. V. Metcalf, L. Zhang, L. Pini, G. L. Shulman, and M. Corbetta, Spontaneous activity patterns in human motor cortex replay evoked activity patterns for hand movements, Scientific Reports 12, 16867 (2022).

[22] J. N. MacLean, B. O. Watson, G. B. Aaron, and R. Yuste, Internal dynamics determine the cortical response to thalamic stimulation, Neuron 48, 811 (2005).

[23] V. Pasquale, S. Martinoia, and M. Chiappalone, Stimulation triggers endogenous activity patterns in cultured cortical networks, Scientific reports 7, 9080 (2017).

[24] M. Angiolelli, S. Scarpetta, P. Sorrentino, E. Troisi Lopez, M. Quarantelli, C. Granata, G. Sorrentino, V. Palmieri, G. Messuti, M. Stefano, et al., Role of criticality in the structure-function relationship in the human brain, Physical Review Research 7, 043153 (2025).

[25] A. P. Vaz, J. Wittig, John H., S. K. Inati, and K. A. Zaghloul, Replay of cortical spiking sequences during human memory retrieval, Science 367, 1131 (2020).

[26] X. Jiang, I. Shamie, W. K. Doyle, D. Friedman, P. Dugan, O. Devinsky, E. Eskandar, S. S. Cash, T. Thesen, and E. Halgren, Replay of large-scale spatio-temporal patterns from waking during subsequent NREM sleep in human cortex, Scientific Reports 7, 17380 (2017).

[27] Q. Huang, Z. Xiao, Q. Yu, Y. Luo, J. Xu, Y. Qu, R. J. Dolan, T. Behrens, and Y. Liu, Replay-triggered brain-wide activation in humans, Nature Communications 15, 7185 (2024).

[28] Z. S. Chen and M. A. Wilson, How our understanding of memory replay evolves, Journal of Neurophysiology 129, 552 (2023).

[29] M. A. Montemurro, M. J. Rasch, Y. Murayama, N. K. Logothetis, and S. Panzeri, Phase-of-firing coding of natural visual stimuli in primary visual cortex, Current Biology 18, 375 (2008).

[30] Kayser, M. A. Montemurro, N. K. Logothetis, and S. Panzeri, Spike-phase coding boosts and stabilizes information carried by spatial and temporal spike patterns, Neuron 61, 597 (2009).

[31] M. Siegel, M. R. Warden, and E. K. Miller, Phase-dependent neuronal coding of objects in short-term memory, Proceedings of the National Academy of Sciences 106, 21341 (2009).

[32] E. Bullmore and O. Sporns, The economy of brain network organization, Nature reviews neuroscience 13, 336 (2012).

[33] S. Scarpetta and A. De Candia, Information capacity of a network of spiking neurons, Physica A: Statistical Mechanics and its Applications 545, 123681 (2020).

[34] H. Markram, J. Lübke, M. Frotscher, and B. Sakmann, Regulation of synaptic efficacy by coincidence of postsynaptic aps and epsps, Science 275, 213 (1997).

[35] G.-q. Bi and M.-m. Poo, Synaptic modifications in cultured hippocampal neurons: dependence on spike timing, synaptic strength, and postsynaptic cell type, Journal of neuroscience 18, 10464 (1998).

[36] L. F. Abbott and S. B. Nelson, Synaptic plasticity: Taming the beast, Nature Neuroscience 3, 1178 (2000).

[37] H. Markram, W. Gerstner, and P. J. Sjöström, Spike-timing-dependent plasticity: A comprehensive overview, Frontiers in Synaptic Neuroscience 4, 10.3389/fnsyn.2012.00002 (2012).

[38] W. Gerstner, R. Kempter, J. L. Van Hemmen, and H. Wagner, A neuronal learning rule for sub-millisecond temporal coding, Nature 383, 76 (1996).

[39] S. Song, K. D. Miller, and L. F. Abbott, Competitive hebbian learning through spike-timing-dependent synaptic plasticity, Nature Neuroscience 3, 919 (2000).

[40] W. Senn, H. Markram, and M. Tsodyks, An algorithm for modifying neurotransmitter release probability based on pre-and postsynaptic spike timing, Neural computation 13, 35 (2001).

[41] W. Gerstner and W. M. Kistler, Mathematical formulations of hebbian learning, Biological cybernetics 87, 404 (2002).

[42] H. D. Abarbanel, R. Huerta, and M. Rabinovich, Dynamical model of long-term synaptic plasticity, Proceedings of the National Academy of Sciences 99, 10132 (2002).

[43] S. Scarpetta, Z. Li, and J. A. Hertz, Hebbian imprinting and retrieval in oscillatory neural networks, Neural Computation 14, 2371 (2002).

[44] C. Meisel and T. Gross, Adaptive self-organization in a realistic neural network model, Physical Review E—Statistical, Nonlinear, and Soft Matter Physics 80, 061917 (2009).

[45] J.-F. Zhou, W.-J. Yuan, D. Chen, B.-H. Wang, Z. Zhou, S. Boccaletti, and Z. Wang, Synaptic modifications driven by spike-timing-dependent plasticity in weakly coupled bursting neurons, Physical Review E 99, 032419 (2019).

[46] M. Gillett, U. Pereira, and N. Brunel, Characteristics of sequential activity in networks with temporally asymmetric hebbian learning, Proceedings of the National Academy of Sciences of the United States of America 117, 29948 (2020).

[47] M. Abeles, Corticonics: Neural circuits of the cerebral cortex (Cambridge University Press, 1991).

[48] S. Scarpetta, A. De Candia, and F. Giacco, Storage of phase-coded patterns via stdp in fully-connected and sparse network: a study of the network capacity, Frontiers in synaptic neuroscience 2, 32 (2010).

[49] S. Scarpetta, F. Giacco, and A. de Candia, Storage capacity of phase-coded patterns in sparse neural networks, Europhysics Letters 95, 28006 (2011).

[50] L. Minati, S. Scarpetta, M. Andelic, P. A. Valdes-Sosa, L. Ricci, and A. de Candia, First-and second-order phase transitions in electronic excitable units and neural dynamics under global inhibitory feedback, Chaos, Solitons & Fractals 182, 114701 (2024).

[51] W. Gerstner and W. M. Kistler, Spiking Neuron Models: Single Neurons, Populations, Plasticity (Cambridge University Press, 2002).

[52] W. Gerstner, W. M. Kistler, R. Naud, and L. Paninski, Neuronal Dynamics: From Single Neurons to Networks and Models of Cognition (Cambridge University Press, 2014).

[53] S. Scarpetta and A. de Candia, Neural avalanches at the critical point between replay and non-replay of spatiotemporal patterns, PLoS One 8, e64162 (2013).

[54] J.-P. Lachaux, E. Rodriguez, J. Martinerie, and F. J. Varela, Measuring phase synchrony in brain signals, Human brain mapping 8, 194 (1999).

[55] L. Marple, Computing the discrete-time” analytic” signal via fft, IEEE Transactions on signal processing 47, 2600 (1999).

[56] S. Scarpetta, F. Giacco, F. Lombardi, and A. De Candia, Effects of poisson noise in a if model with stdp and spontaneous replay of periodic spatiotemporal patterns, in absence of cue stimulation, Biosystems 112, 258 (2013).

[57] S. Scarpetta and A. De Candia, Alternation of up and down states at a dynamical phase-transition of a neural network with spatiotemporal attractors, Frontiers in systems neuroscience 8, 88 (2014).

[58] J. M. Beggs and D. Plenz, Neuronal avalanches in neocortical circuits, Journal of Neuroscience 23, 11167 (2003).

[59] O. Shriki et al., Neuronal avalanches in the resting meg of the human brain, Journal of Neuroscience 33, 7079 (2013).

[60] W. L. Shew and D. Plenz, The functional benefits of criticality in the cortex, The Neuroscientist 19, 88 (2013).

[61] P. Massobrio, L. de Arcangelis, V. Pasquale, H. J. Jensen, and D. Plenz, Criticality as a signature of healthy neural systems (2015).

[62] G. Ódor and B. De Simoni, Heterogeneous excitable systems exhibit griffiths phases below hybrid phase transitions, Physical Review Research 3, 013106 (2021).

[63] G. Ódor, I. Papp, S. Deng, and J. Kelling, Synchronization transitions on connectome graphs with external force, Frontiers in Physics 11, 1150246 (2023).

[64] K. B. Hengen and W. L. Shew, Is criticality a unified set-point of brain function?, Neuron, Volume 113, Issue 16 (2025).

[65] K. Srinivasan, T. L. Ribeiro, P. Kells, and D. Plenz, The recovery of parabolic avalanches in spatially subsampled neuronal networks at criticality, Scientific Reports 14, 19329 (2024).

[66] A. Zhigalov, G. Arnulfo, L. Nobili, S. Palva, and J. M. Palva, Modular co-organization of functional connectivity and scale-free dynamics in the human brain, Network Neuroscience 1, 143 (2017).

[67] H. Yamamoto, F. P. Spitzner, T. Takemuro, V. Buendía, H. Murota, C. Morante, T. Konno, S. Sato, A. Hirano-Iwata, A. Levina, et al., Modular architecture facilitates noise-driven control of synchrony in neuronal networks, Science advances 9, eade1755 (2023).

[68] V. Myrov, A. Suleimanova, S. Knapič, P. Partanen, M. Vesterinen, W. Liu, S. Palva, and J. M. Palva, Hierarchical whole-brain modeling of critical synchronization dynamics in human brains, bioRxiv, 2024 (2024).

[69] M. T. Cirunay, R. C. Batac, and G. Ódor, Learning and criticality in a self-organizing model of connectome growth, Scientific Reports 15, 31890 (2025).

[70] P. Del Giudice, S. Fusi, and M. Mattia, Modelling the formation of working memory with networks of integrate-and-fire neurons connected by plastic synapses, Journal of Physiology, Paris 97, 659 (2003).

[71] M. Lengyel, J. Kwag, O. Paulsen, and P. Dayan, Matching storage and recall: hippocampal spike timing-dependent plasticity and phase response curves, Nature Neuroscience 8, 1677 (2005).

[72] J. K. Liu and D. V. Buonomano, Embedding multiple trajectories in simulated recurrent neural networks in a self-organizing manner, Journal of Neuroscience 29, 13172 (2009).

[73] T. Masquelier, E. Hugues, G. Deco, and S. J. Thorpe, Oscillations, phase-of-firing coding, and spike timing-dependent plasticity: an efficient learning scheme, Journal of Neuroscience 29, 13484 (2009).

[74] M. Gilson, A. Burkitt, and J. L. van Hemmen, Stdp in recurrent neuronal networks, Frontiers in Computational Neuroscience 4, 23 (2010).

[75] R. Fiete, W. Senn, C. Z. Wang, and R. H. Hahnloser, Spike-time-dependent plasticity and heterosynaptic competition organize networks to produce long scale-free sequences of neural activity, Neuron 65, 563 (2010).

[76] P. Zheng and J. Triesch, Robust development of synfire chains from multiple plasticity mechanisms, Frontiers in Computational Neuroscience 8, 66 (2014).

[77] A. Litwin-Kumar and B. Doiron, Formation and maintenance of neuronal assemblies through synaptic plasticity, Nature Communications 5, 5319 (2014).

[78] F. Zenke, E. J. Agnes, and W. Gerstner, Diverse synaptic plasticity mechanisms orchestrated to form and retrieve memories in spiking neural networks, Nature Communications 6, 6922 (2015).

[79] F. Weissenberger, F. Meier, J. Lengler, H. Einarsson, and A. Steger, Long synfire chains emerge by spike-timing dependent plasticity modulated by population activity, International Journal of Neural Systems 27, 1750044 (2017).

[80] G. K. Ocker and B. Doiron, Training and spontaneous reinforcement of neuronal assemblies by spike timing plasticity, Cerebral Cortex 29, 937 (2019).

[81] U. Pereira and N. Brunel, Unsupervised learning of persistent and sequential activity, Frontiers in Computational Neuroscience 13, 97 (2020).

[82] M. Gillett, U. Pereira, and N. Brunel, Characteristics of sequential activity in networks with temporally asymmetric hebbian learning, Proceedings of the National Academy of Sciences 117, 29948 (2020).

[83] R. Bergoin, A. Torcini, G. Deco, M. Quoy, and G. Zamora-López, Emergence and maintenance of modularity in neural networks with hebbian and anti-hebbian inhibitory stdp, PLOS Computational Biology 21, e1012973 (2025).

[84] S. Panzeri, M. Moroni, H. Safaai, and C. D. Harvey, The structures and functions of correlations in neural population codes, Nature Reviews Neuroscience 23, 551 (2022).

[85] Russo, N. Becker, A. P. F. Domanski, T. Howe, K. Freud, D. Durstewitz, and M. W. Jones, Integration of rate and phase codes by hippocampal cell-assemblies supports flexible encoding of spatiotemporal context, Nature Communications 15, 8880 (2024).

[86] Londei, F. Ceccarelli, G. Arena, L. Ferrucci, E. Russo, E. Brunamonti, and A. Genovesio, Out of the single-neuron straitjacket: Neurons within assemblies change selectivity and their reconfiguration underlies dynamic coding, The Journal of Physiology 603, 4063 (2025).

[87] S. Panzeri, J. H. Macke, J. Gross, and C. Kayser, Neural population coding: combining insights from microscopic and mass signals, Trends in Cognitive Sciences 19, 162 (2015).

[88] M. Shine, Neuromodulatory influences on integration and segregation in the brain, Trends in cognitive sciences 23, 572 (2019).

[89] Barzon, M. Allegra, M. H. Aarabi, L. Pini, M. De Domenico, M. Corbetta, and S. Suweis, Structural connectome dimension shapes brain dynamics in health and disease, bioRxiv 10.1101/2025.06.30.662336 (2025), preprint.

[90] P. Sorrentino, G. Rabuffo, F. Baselice, E. Troisi Lopez, M. Liparoti, M. Quarantelli, G. Sorrentino, C. Bernard, and V. Jirsa, Dynamical interactions reconfigure the gradient of cortical timescales, Network Neuroscience 7, 73 (2023).

[91] Varela, J.-P. Lachaux, E. Rodriguez, and J. Martinerie, The brainweb: phase synchronization and large-scale integration, Nature reviews neuroscience 2, 229 (2001).

[92] M. A. Long, D. Z. Jin, and M. S. Fee, Support for a synaptic chain model of neuronal sequence generation, Nature 468, 394 (2010).

[93] D. J. Amit and D. J. Amit, Modeling brain function: The world of attractor neural networks (Cambridge university press, 1989).

[94] N. Brunel, Is cortical connectivity optimized for storing information?, Nature neuroscience 19, 749 (2016).

