## Supplementary figures for "Modularity-dependent storage of dynamic spiking patterns: bridging micro- and mesoscopic representations"

### Supplementary Material

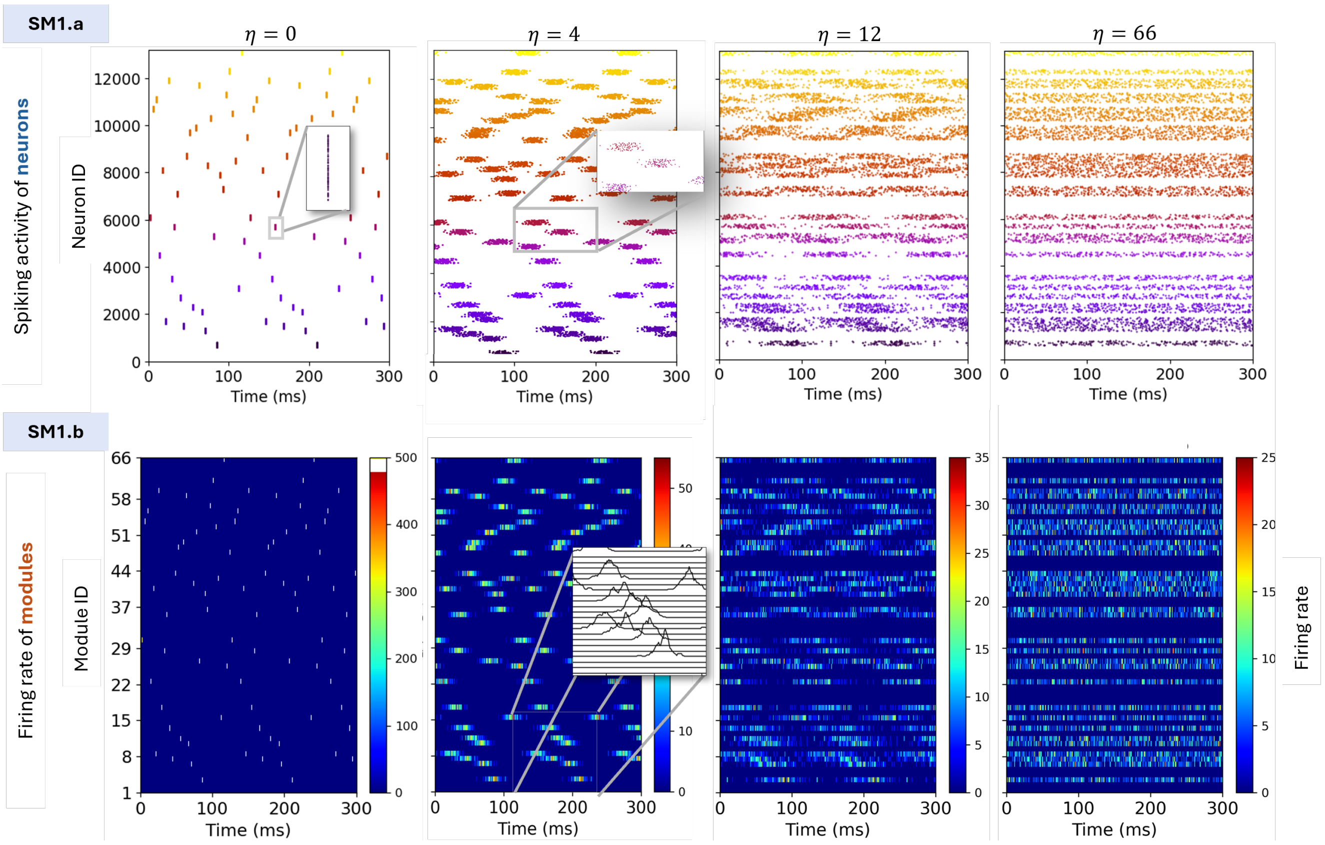


Figure SM1: **Encoding of the second modular spatio-temporal pattern during learning.** Each stored periodic pattern is an ordered sequence of $G \leq S$ active modules, with overlap between modules regulated by $\eta$, and precise spike timing for the $K \leq Z$ non-silent neurons. Activity at increasing $\eta$ values (0,4,12,66) is shown with $G = 33, K = 100$ for one specific pattern. Each panel in (a) shows a raster plot of the neurons involved into the pattern sorted according to the pattern phase inside each module. At $\eta = 0$, neurons within each module are highly synchronised, firing almost simultaneously, while each module is engaged sequentially. As $\eta$ increases, spike timing between groups becomes less distinct. In other words, neurons belonging to a given module begin to fire before the activity of the preceding module has entirely ceased, leading to increasing overlap between the activity of different modules. For high $\eta$, firing becomes broadly distributed with substantial temporal overlap between neuronal modules, reflecting a more homogeneous activation across the network. This behaviour is also visible in panels (b), showing the firing rate of each module, computed using 1 ms bins and expressed in Hz per neuron.

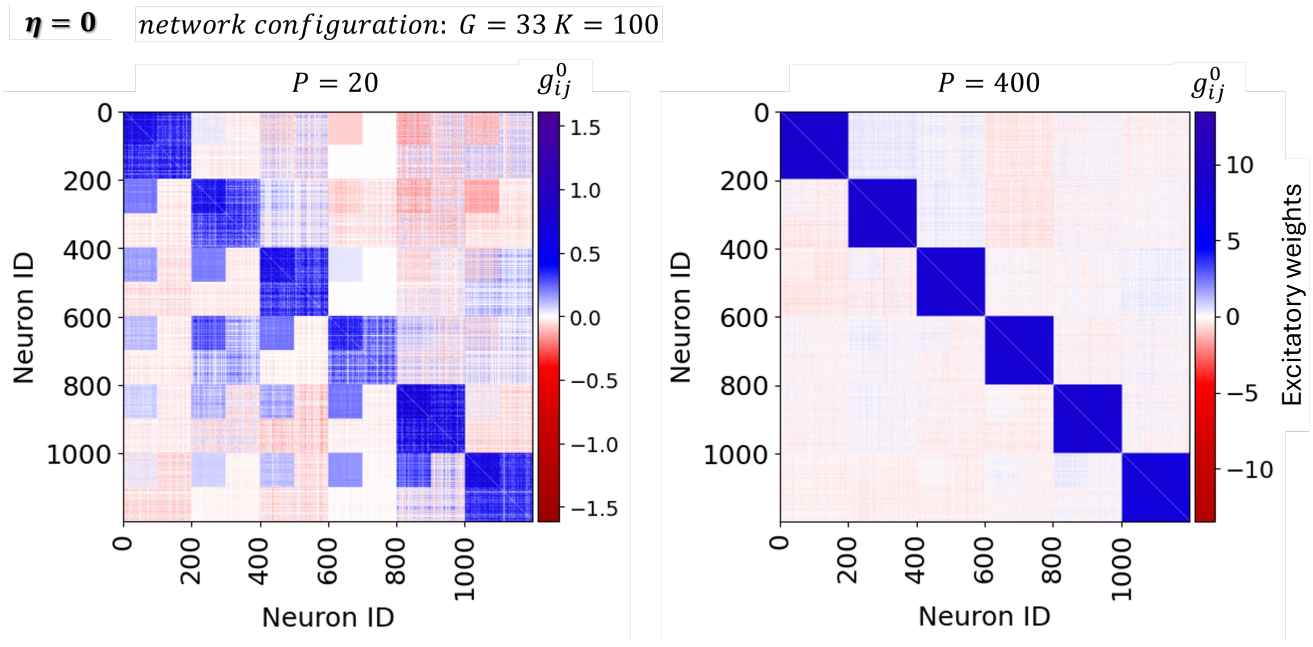


Figure SM2: **Effect of $\eta = 0$ on the network connectivity for one configuration of the network and different numbers of patterns stored into the network.** Connectivity matrix $(g_{ij}^{\eta})$ between individual neurons, when $P = 20$ (left) and $P = 400$ (right) patterns are stored with $\eta = 0$, in a network with $G = 33, K = 100$. Only the first 1200 neurons (corresponding to the first 6 modules) are shown for readability. For $P = 20$, it is more clearly visible that even at $\eta = 0$ (as also clearly occurs for larger $\eta$), the STDP window is able to concatenate multiple modules.

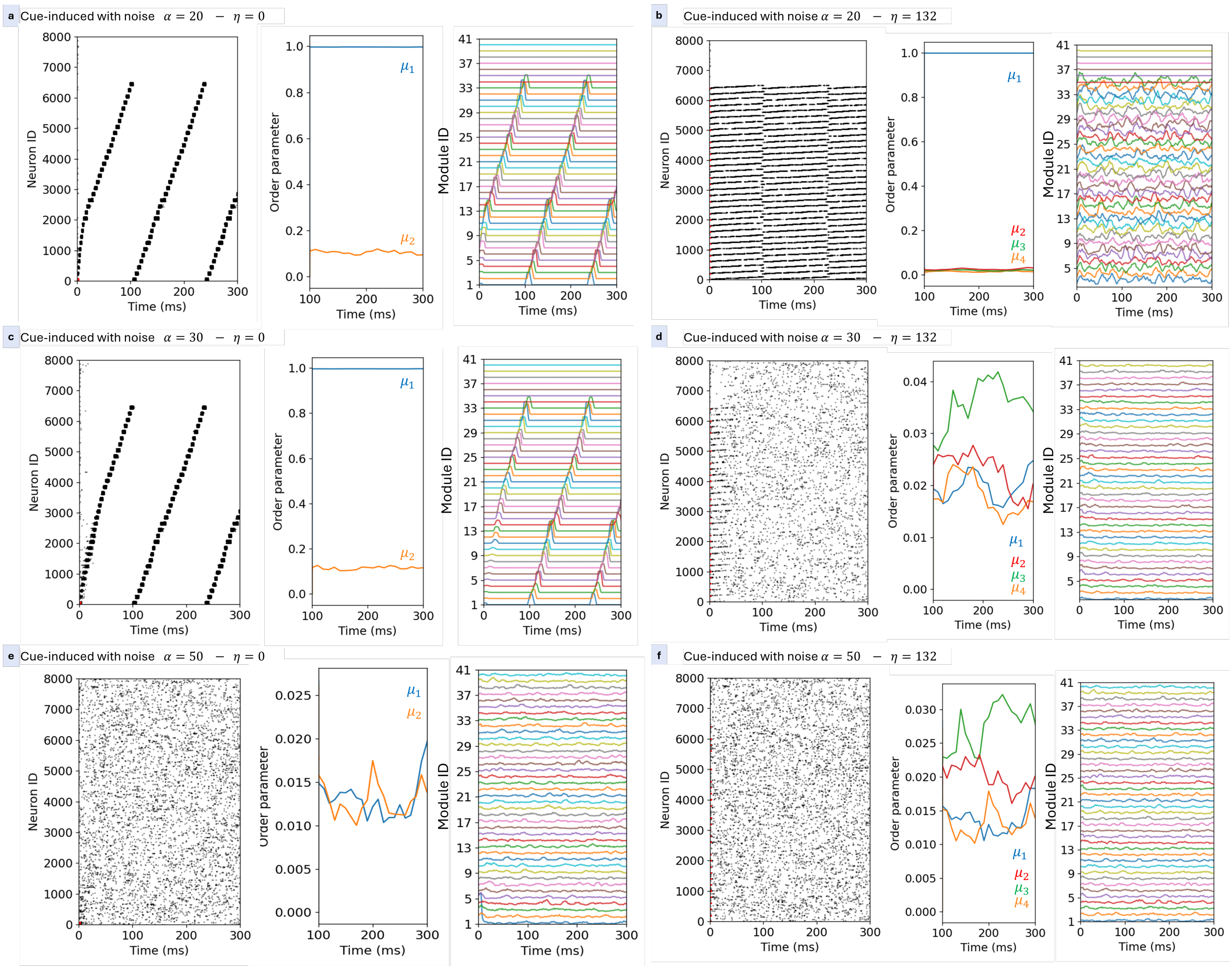


Figure SM3: Effect of noise on cue-induced pattern recall. In each panel, raster plots, order parameters, and module firing rates are shown when both cue and noise (as defined in Eq. 12) are applied as input to the network. The same network parameters as in Fig.4 panels a,b ($G = 33, K = 100, E_0 = 5.5, I_0 = 0.6$) are used. Panels correspond to different values of the parameter $\alpha$ (rows: $\alpha = 20, 30, 50$), representing increasing noise intensity, and two values of the module co-activation index ($\eta$) (columns: $\eta = 0$ and $\eta = 132$). In all cases, recall is initiated by a small cue ($H = 75$ spikes) to retrieve the first stored pattern. For $\alpha = 20$, replay remains robust, the overlap is high, and the raster plots closely match those shown in Fig. 4a,b without noise. At higher noise levels, replay becomes progressively corrupted, and at the largest noise intensity, it is completely disrupted.

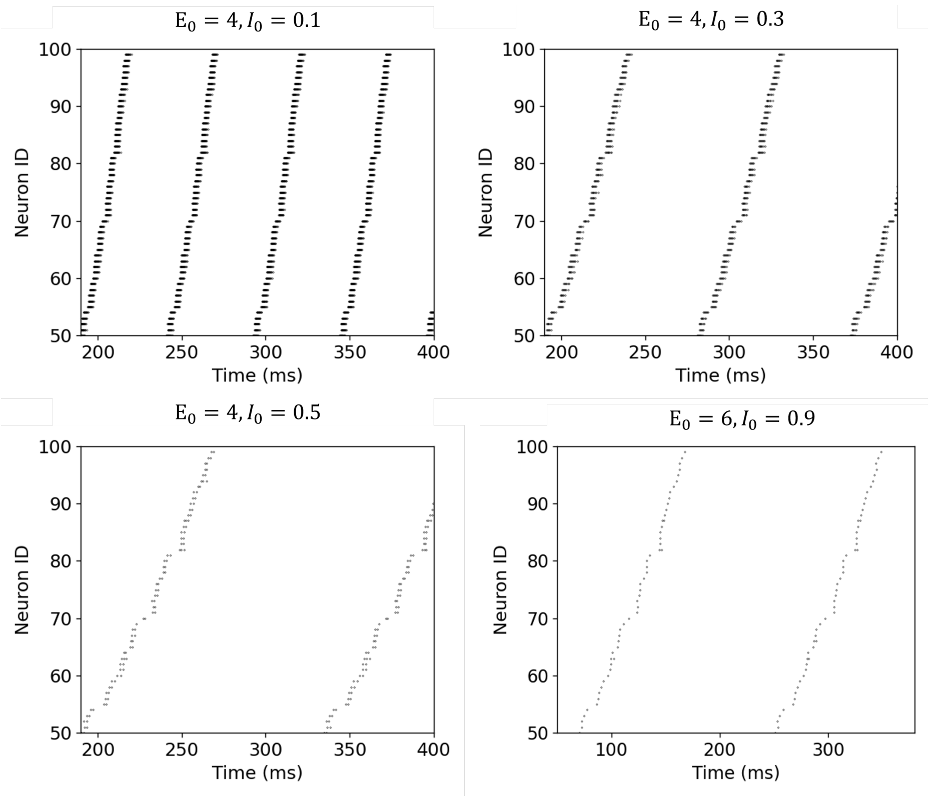


Figure SM4: Excitation–inhibition balance ($E_0/I_0$) modulates period of replay and the number of spikes per cycle, preserving the phase relationship among units. Recall of the pattern $\mu = 1$ for the configuration network with $G = 33, K = 100, P = 10, \eta = 20$. Different values of $E_0$ and $I_0$ are shown. Increasing global inhibition$I_0$ at fixed $E_0$ reduces the number of spikes per neuron per cycle without disrupting phase-coded replay. Only 50 neurons are shown for clarity.

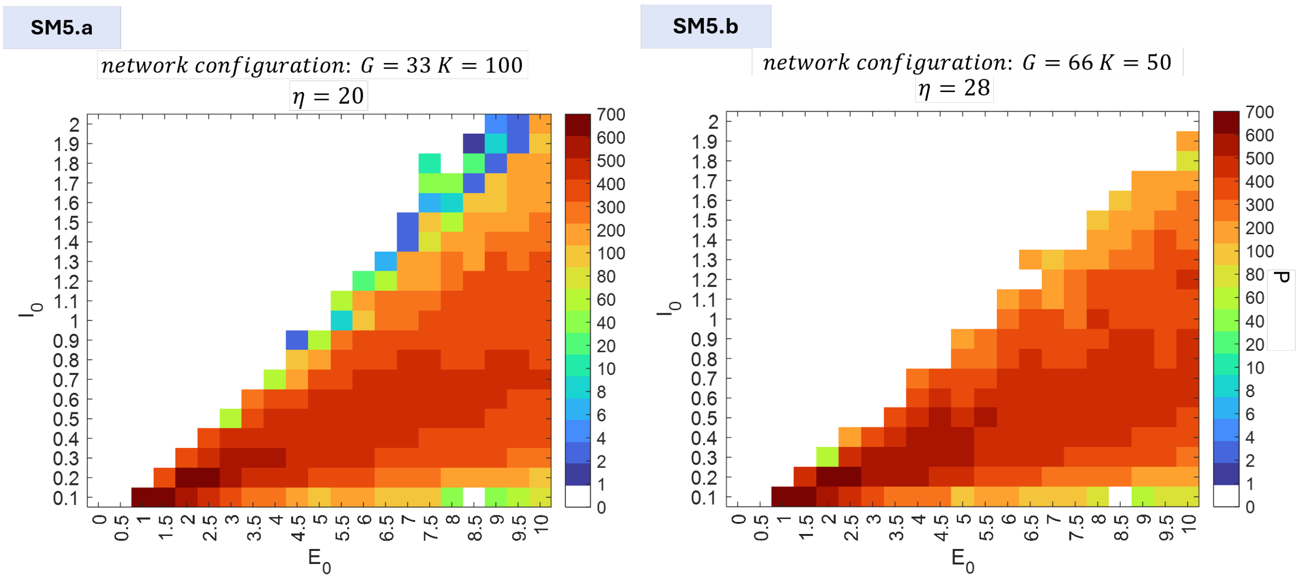


Figure SM5: **Maximum number of patterns as a function of $E_0$ and $I_0$ at a crucial $\eta$ values**. Panels (a) and (b) show the maximum number of patterns $P_{\max}$ as a function of the parameters $(E_0, I_0)$ at $\eta = 20$ and $\eta = 28$ for the $(G = 33, K = 100)$ and $(G = 66, K = 50)$ configuration, respectively. These $\eta$ values correspond to the smallest $\eta$ at which the network reaches its maximum storage capacity.
